# Epigenetic remodeling via HDAC6 inhibition amplifies anti-tumoral immune responses in myeloid leukemia cells

**DOI:** 10.1101/2025.05.30.654805

**Authors:** Julian Schliehe-Diecks, Jia-Wey Tu, Pawel Stachura, Katerina Schaal, Marie Kemkes, Eleni Vasileiou, Nadine Rüchel, Danielle Brandes, Melina Vogt, Thomas Lenz, Adarsh Nair, Stefanie Scheu, Pilar M. Dominguez, Agata Pastorczak, Karin Nebral, Kai Stühler, Ute Fischer, Aleksandra A. Pandyra, Arndt Borkhardt, Sanil Bhatia

## Abstract

Histone deacetylase 6 (HDAC6) has emerged as a promising therapeutic target in cancer due to its immunomodulatory effects. While its prognostic significance remains debated, we demonstrate that HDAC6 loss significantly impairs myeloid leukemia progression in vivo, despite having no functional impact on leukemia cell proliferation in vitro. Global proteome and secretome profiling of HDAC6-knockout (KO) cells revealed upregulation of several immune-related modulators, including RNase T2, a tumor suppressor known to modulate the tumor microenvironment. Notably, RNase T2 upregulation upon HDAC6 loss was restricted to myeloid but not B-ALL cells. Moreover, pharmacological inhibition of HDAC6 recapitulated this phenotype, leading to RNase T2 upregulation in myeloid leukemia cells. ATAC-seq revealed increased chromatin accessibility of RNase T2 following HDAC6 loss, highlighting a functionally epigenetic regulatory contribution. Further functional assays conducted in an immunocompetent setting both ex vivo and in vivo demonstrated that HDAC6 inhibition sensitized murine myeloid leukemia cells to broad CD8^+^ T cell activation as evidenced by increased TNFα and CD107a expression. Consistently, in a syngeneic model, HDAC6 inhibition restricted growth myeloid leukemia cells. Moreover, an extended drug screening analysis identified Cytarabine and Clofarabine as significantly synergizing with HDAC6 inhibitor (Ricolinostat) in myeloid leukemia cell lines and in patient derived xenograft (PDX) cells, while showing limited synergy in lymphoid leukemia cell lines, PDX or healthy control cells. These findings suggest that HDAC6 represents a promising therapeutic target in myeloid lineage derived leukemia cells by simultaneously enhancing immune activation and increasing chemosensitivity.

## Introduction

Aberrant epigenetic modifications, including histone deacetylation, play a key role in the leukemogenesis of acute myeloid leukemia (AML) and contribute to chemotherapy resistance by silencing tumor suppressor genes regulating chemosensitivity.^1^ Dysregulation of HDACs has been linked to impaired hematopoietic differentiation, cell cycle defects, DNA damage accumulation, and reduced cell viability.^2^ Although mutations in HDACs are uncommon in myeloid malignancies, dysregulated expression and aberrant recruitment by leukemia-associated fusion proteins have been reported.^3,4^ HDAC inhibitors (HDACi) have recently emerged as a promising class of therapeutics for various malignancies, including AML.^5–7^ However, the approved HDACi for treating hematological malignancies are predominantly pan-HDACi, which broadly inhibit multiple HDAC isoforms involved in essential cellular functions.^8,9^ This lack of specificity leads to a range of adverse effects, including cardiac toxicity and severe gastrointestinal disturbances, some of which can be life-threatening.^10–12^ These toxicities often require the discontinuation of HDACi-based therapies to prevent severe complications. Additionally, in clinical practice, HDACi are rarely used as monotherapy and are typically combined with other therapeutic agents, adding complexity to treatment regimens.^13,14^ Selective targeting of specific HDAC isoforms may offer a promising approach to restricting their activity to particular cellular processes, potentially reducing toxicity and enhancing therapeutic tolerability.

Among the various HDAC isoforms, HDAC6 has emerged as a particularly compelling therapeutic target due to its distinct biological roles and relevance in oncogenesis.^15^ HDAC6 is primarily localized in the cytoplasm and regulates a range of cellular functions through substrate deacetylation.^16–19^ However, HDAC6-deficient mice remain viable and develop normally, supporting its potential as a safe therapeutic target.^20^ HDAC6 is found overexpressed across various hematologic malignancies.^21–23^ Pharmacological HDAC6 inhibition has been shown to target leukemic stem cells (LSCs) in CML.^22,24,25^ A recent study from a zebrafish model demonstrates that inhibition of HDAC6 suppresses LSC expansion and reduces the proliferation of AML cell lines,^26^ underscoring its potential as a therapeutic target across myeloid malignancies.^27^ However, the prognostic relevance of HDAC6 as a therapeutic target remains debated, as HDAC6 inhibition alone has been reported to exert minimal or no cytotoxic effects on leukemia cells in ex vivo studies.^28–30^

Through a database-driven approach, we identified HDAC6 as a promising candidate in AML. Using HDAC6-Knockout (KO) models and employing multi-omics characterization we identified RNase T2, a known activator of pro-inflammatory signaling, to be upregulated upon HDAC6-KO. Further, our ex vivo and in vivo studies demonstrated that HDAC6 inhibition enhanced broad CD8^+^ T cell activation and therefore anti-leukemic responses. Beyond its immunomodulatory role, we found that HDAC6 inhibition sensitized myeloid leukemia cells to standard chemotherapeutics. These findings highlight the potential of HDAC6 as a target in myeloid leukemia, underscoring the need for further investigation into its clinical applicability.

## Methods

### Cell Culture

Leukemia cell lines were obtained from the DSMZ (Braunschweig, Germany) and cultured according to their recommendations. RPMI1640 GlutaMAX (Gibco) was supplemented with 10-20% fetal bovine serum and 1% penicillin/streptomycin (Sigma-Aldrich). Cell lines were regularly authenticated by short tandem repeat profiling and tested for mycoplasma.

### CRISPR-Cas9 mediated knockout (KO) or shRNA-mediated knockdown (KD) of HDAC6

HDAC6-KO models were generated either through lentiviral transduction or using nucleofection of CAS9-GFP (IDT).^31^ For monoclonal selection, cells were plated in semi-solid MethoCult media (Stemcell Technologies). See supplementary methods for sequences and more details.

### Western blotting

Western blotting were performed as previously described.^31^ For the list of antibodies and their respective concentrations, refer to the supplementary methods.

### Proliferation, cell cycle and apoptosis assays

Proliferation (via Trypan Blue staining), cell cycle (via Nicoletti method) and apoptosis assay (via measuring Caspase 3/7 activity) were performed as described earlier.^31^

### Colony forming unit (CFU) assay

Leukemia were seeded in a methylcellulose-based medium. After seven days, colonies (n=3) were counted and imaged, following previously described protocols.^32^

### Drug screening

Ex vivo high throughput drug screening was performed as described earlier.^31,33^ The compound selection comprised FDA/EMA-approved chemotherapeutics and targeted agents commonly used in leukemia treatment protocols, along with inhibitors in various stages of clinical trials. A detailed list of drugs is provided in supplementary Table 1. ZIP scores for the matrix synergy approach were calculated via the *Synergyfinder* R package.

### qPCR

Total RNA was extracted using QIAzol and purified with the Maxwell® RSC system. cDNA was synthesized from 2 µg RNA using the QuantiTect kit, and qPCR was performed on a Bio-Rad system with B2M and GAPDH as internal controls. For primer sequences and additional details, refer to supplementary methods.

### Fluorescence Microscopy

Cells were seeded on poly-D-lysine–coated coverslips, fixed, permeabilized, and stained with primary/secondary antibodies and DAPI. Imaging was performed with a Zeiss Axio Observer/Apotome 3, and signal quantification was done using ImageJ. For more details, refer to supplementary methods.

### Murine NSG transplantation

Prior to the injection K562 and MV4-11 cells were transduced with a luciferase reporter plasmid to measure the leukemic burden via the in vivo imaging system IVIS (PerkinElmer).^31^ Cells were intravenously injected into 6 to 8 weeks old immunodeficient NSG mice (Jackson laboratory, #005557). For patient-derived xenograft (PDX) generation, leukemia cells from peripheral blood or bone marrow were injected into NSG mice (see Table S2 for patient details), with engraftment monitored via FACS analysis of blood collected through orbital bleeding.^31,34^ Patient material was obtained with informed consent in accordance with the Declaration of Helsinki and approved by the local ethics committees. All animal experiments were conducted in accordance with the regulatory guidelines of the official committee at LANUV (NRW), under the authorization of the animal research institute (ZETT) at the Heinrich Heine University Düsseldorf.

### Murine C57BL/6J experiments

Wildtype C57BL/6J (Charles River) mice was utilized to establish murine C1498 AML (ATCC) derived syngeneic mouse model.^35^ C57BL/6J wild type mice were engrafted with (0.5x10^6 cells/mouse) luciferase reporter gene transduced C1498 murine AML cell line at 6-7 weeks of age. Monitoring of the leukemic burden and treatment with Ricolinostat followed the respective schedule. For more details, refer to supplementary methods.

### T Cell Co-Culture Assay

C57BL/6J mice were infected with the Lymphocytic Choriomeningitis Virus (LCMV) strain, which induces a strong effector CD8⁺ T cell response and is efficiently cleared in wild-type mice shortly after infection.^35,36^ LCMV-primed splenic T cells were isolated after two weeks post infection. C1498 cells were pre-treated with Ricolinostat before co-culture with T cells. Supernatant-only and direct treatment conditions were also included. After 24 hours, cells were analyzed by flow cytometry for viability and cytokine expression. For more details, refer to supplementary methods.

### NK Cell Killing Assay

PBMCs from healthy donors were cultured with IL-2 and IL-15. CFSE-labeled target cells were co-incubated with PBMC-derived NK cells for 4 hours, followed by flow cytometric analysis to assess NK cell-mediated cytotoxicity. For more details, refer to supplementary methods.

### RNA-sequencing (RNA-seq), mass spectrometery (MS) based proteome and secretome anaylsis

RNA-seq and quantitative MS based proteome and secretome analysis was performed as previously described.^31^ For sample preparation, cultures were maintained at a medium density for several days to harmonize viability and growth rates.

### ATAC-seq

The library preparation, sequencing and initial consensus sequencing was performed according to the protocol of Active Motif. For more details, refer to supplementary methods.

### Data mining

The HDAC6 dependency data was extracted from the Achilles gene dependency project of the DepMap (22Q1) database and visualized via complex heatmap.^37^ The expression data of pediatric patients was extracted from the PeCan platform of St. Jude.^38^ Expression was either clustered by leukemia subtype or separated by the median of HDAC6 expression. The survival analysis was performed using the Survival Genie 2 platform^39^, utilizing data generated by the Therapeutically Applicable Research to Generate Effective Treatments (TARGET) initiative (https://www.cancer.gov/ccg/research/genome-sequencing/target), phs000218.^40^ The data used for this analysis are available through the Genomic Data Commons (https://portal.gdc.cancer.gov). Samples were clustered by their optimal cutting point algorithm. Mutation clustered survival analysis was performed via Kaplan-Meier-plotter^41^ which utilizes the TCGA-LAML data set. Patients with IDH1, NRAS or FLT3 mutation were split by a percentile-based best cut-off selection.

Pathway activity scores were inferred using Pathifier,^42^ and transcription factor activity was estimated using VIPER with the DoRothEA regulon,^43^ based on gene expression data from AML and CML cell lines in the DepMap (25Q2) dataset. The resulting activity profiles were then correlated with ZIP synergy scores of the Ricolinostat and Cytarabine or Clofarabine combination using both Pearson and Spearman correlation coefficients.

### Data Availability

RNA-seq and ATAC-seq data have been deposited in the NCBI GEO database under accession ID: GSE297364 and GSE297366. Mass spectrometry-based proteomics and secretomics data are available via the ProteomeXchange Consortium through the PRIDE repository with dataset identifier PXD041871.

## Results

### HDAC6 ablation impairs in vivo growth of myeloid leukemia cells

We data-mined clinical data using TARGET and TCGA-LAML datasets, and identified prognostic implications. High HDAC6 expression significantly correlated with poorer survival in AML, whereas in ALL, elevated HDAC6 levels were associated with improved survival outcomes (Figure S1A-H). Further, the analysis of over 1,500 leukemia patient samples (from the St Jude. PeCan database) revealed a more heterogeneous HDAC6 expression pattern in AML compared to B-ALL and T-ALL (coefficient of variance or CV_AML_=0.70 vs CV_B-ALL_=0.48 or CV_T-ALL_=0.39) (Figure S1I). These findings contrast with earlier reports suggesting that selective HDAC6 inhibition has minimal or no cytotoxic effects on the leukemia cells in ex vivo studies.^28–30^ Further, in vitro CRISPR-based dependency analysis (using data from the DepMap portal) also suggest that HDAC6 exhibits the lowest dependency among HDAC family members in leukemia cells (Figure S1J).

Given these discrepancies between in vitro and in vivo HDAC6 inhibition, we employed a CRISPR-Cas9-mediated knockout (KO) approach in myeloid leukemia cells to assess the effects of HDAC6 loss. HDAC6-KO models were established in the (*BCR::ABL1^+^*) K562 CML cells and (*KMT2A-r*) MV4-11 AML cells (Figure 1A-H). Loss of HDAC6 resulted in increased α-tubulin acetylation, confirming functional impairment of HDAC6 in these models (Figure 1A and 1E). First, to evaluate the differences in the cellular growth, a proliferation assays were performed, which revealed no significant differences in doubling time between HDAC6-KO and control cells (Figure 1B and 1F). Similarly, a colony formation assay showed no alterations in clonogenic potential upon HDAC6 loss (Figure 1C and 1G). Cell cycle analysis further demonstrated minimal changes in the distribution of cells across different phases (Figure 1D and 1H). However, we observed an increased susceptibility of K562 HDAC6-KO cells to NK cell-mediated killing in an ex vivo PBMC-derived NK cell killing assay, compared to the respective controls (Figure S1K). Notably, in an in vivo setting, HDAC6-KO (K562 and MV4-11) leukemia models exhibited a significant (*p* < 0.05) reduction in leukemia progression when transplanted into NSG mice compared to controls (Figure 1I-L). This led to a significant (p < 0.01) increase in overall survival in mice injected with HDAC6-KO K562 cells, while no significant differences in survival was observed in the MV4-11 HDAC6-KO model. (Figure S1L-M). This discrepancy in survival outcomes may be attributed to the more aggressive disease kinetics of AML-derived MV4-11 cells relative to CML-derived K562 cells, potentially diminishing the observable survival benefit at later stages despite early and significant reductions in leukemic burden. Taken together, our data demonstrate that despite the lack of effect on in vitro growth, HDAC6 loss inhibits progression of myeloid leukemia cells in vivo.

**Figure 1.**
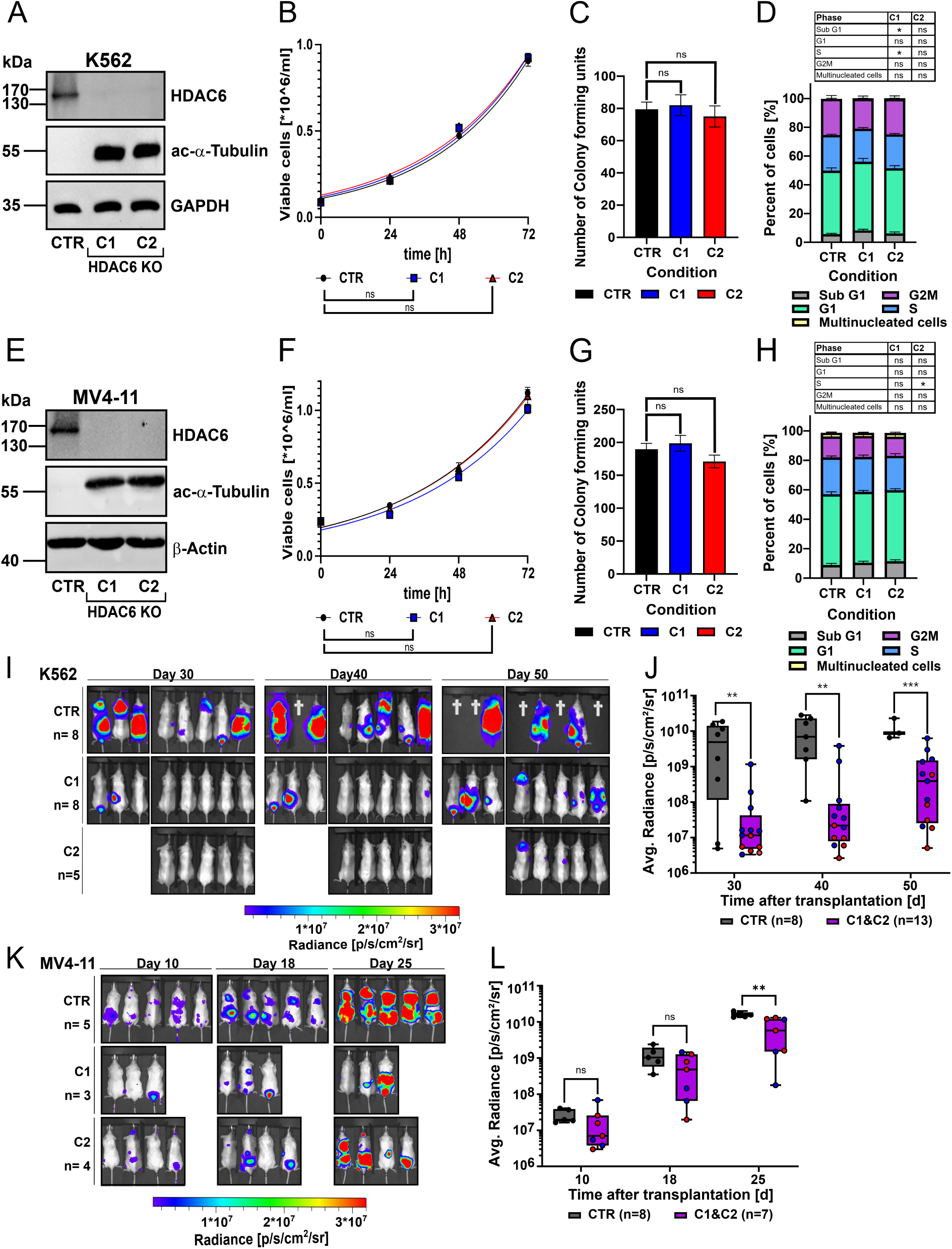
HDAC6 ablation impairs the in vivo growth of myeloid leukemia cells. (A) Representative western blot for HDAC6 and acetylated α-tubulin protein levels in 2 clones (C1 and C2) of K562 HDAC6-knockout (KO) and a non-targeting control (CTR) (n=3). (B) Proliferation curve determined by trypan blue assay of K562 HDAC6-KO. Statistical significance was calculated by comparing the doubling time of the growth curve (unpaired t-test, ns = not significant, n = 3). (C) Colony forming unit (CFU) assay performed by seeding K562 HDAC6-KO cells in a semi-solid medium (unpaired t-test, ns = not significant, n = 3). (D) Bar plot showing cell cycle of K562 HDAC6-KO analysis (unpaired t-test, ns = not significant, n = 3). (E-H) Similarly, analysis was performed in the MV4-11 HDAC6-KO model, including western blot (E), proliferation assay (F), CFU assay (G) and cell cycle analysis (H) (unpaired t-test ns = not significant, *p < 0.05, n = 3). (I-K) *In vivo* growth kinetics of K562 (I) and MV4-11 (K) leukemia cells in NSG mice, monitored at the indicated time points post-injection using bioluminescence imaging. (J-L) Quantification of the average radiance of (I-K) for K562 (J) and MV4-11 (L) models (Mann-Whitney U-test, ns = not significant, *p < 0.05, **p < 0.01, ***p < 0.001).

### Targeting HDAC6 upregulates lysosomal-related protein accumulation

Building on the impaired leukemia progression observed upon HDAC6 loss, we employed quantitative mass spectrometry (MS) to profile global changes in the cellular proteome and secretome (Figure 2A-B). Gene set enrichment analysis from the proteome analysis identified multiple dysregulated functional pathways upon HDAC6 loss (Figure 2C & Figure S2A-B). Among the most significantly (*FDR* < 0.05) enriched functional clusters in the proteomic analysis were lysosome-associated proteins, including LAMP1, LAMP2 and RNAse T2. Secretome analysis further validated an upregulation of lysosomal enzymes, suggesting increased extracellular release of lysosomal content (Figure 2B). Additionally, HDAC6-KO cells exhibited enrichment of pathways related to immune activation, inflammatory responses, MAPK signaling and cellular motility (Figure 2C). Functional enrichment analysis using STRING confirmed significant (*FDR* < 1.E-15) changes in lysosomal and immune-related pathways, including neutrophil degranulation and innate immune system processes, among the overlapping proteins between the proteome and secretome profiles (Figure 2D and Figure S2C-D). Using fluorescence microscopy, a significant (*p* < 0.01) increase in LAMP1 expression was validated across HDAC6-KO generated models, including K562, KCL22 and MV4-11 (Figure 2E and Figure S2E). Next, we aimed to pharmacologically replicate the effects of HDAC6 loss using a HDAC6 inhibitor, Ricolinostat.^44,45^ Ricolinostat treatment also resulted in a concentration dependent increase in the LAMP1 expression in the (*BCR::ABL1*^+^) PDX-CML cells (Figure 2F). Together, these results demonstrate that targeting HDAC6 inhibition impacts lysosomal and immune-related pathways.

**Figure 2.**
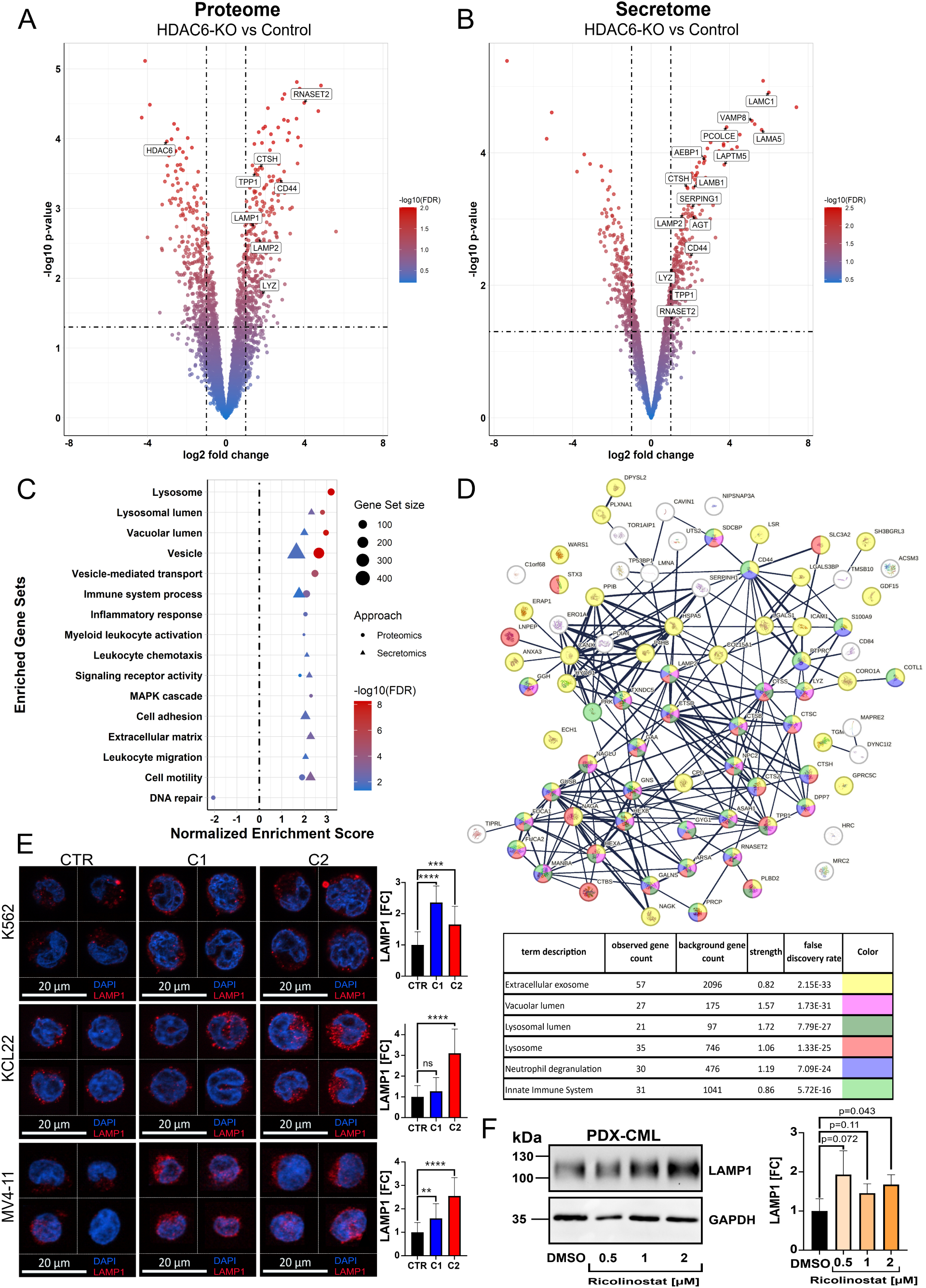
Targeting HDAC6 upregulates lysosome-related protein accumulation. (A-B) Volcano plot depicting the cellular (A) or secreted (B) protein log2 fold change against the negative log_10_(p-value) determined from five independent replicates of K562 HDAC6-KO (C1). Cut off for the log2 fold change was set to >1 while for the -log_10_(p-value) >1.301 (equals <0.05) was used. The color code represents the - log_10_(FDR) (Significance analysis of microarrays method, n= 4-5). (C) Dot plot showing the manually curated enriched gene sets obtained by a *clusterprofiler* analysis for both proteome and secretome results. The FDR cut off for inclusion of protein entries was set to < 0.1 (Kolmogorov-Smirnov test, Benjamini-Hochberg correction). (D) Protein interaction network of the overlapping proteins that were enriched both in the proteomics and secretomics analysis via string. Inclusion criteria was a shared positive direction of enrichment and an FDR < 0.1 (hypergeometric test, Benjamini-Hochberg correction). (E) Fluorescence microscopy analyzing the localization and amount of LAMP1 in K562, KCL22 and MV4-11 HDAC6-KO models. Representative cells are shown while in the right panel the bar graph compares the fold change of fluorescence signal across all analyzed cells (unpaired t-test, ns= not significant, **p < 0.01, ***p < 0.001, ****p < 0.0001, n=20). (F) Western blot analysis of LAMP1 and GAPDH expression of PDX-CML cells treated with Ricolinostat, and in the right panel quantification results normalized to GAPDH (unpaired t-test, n=3).

### Targeting HDAC6 induces the expression of tumor suppressor RNase T2 in myeloid leukemia cells

Next, we selected the lysosomal lumen-associated endoribonuclease RNase T2 as a potential target downstream of HDAC6, based on its well-established role as a tumor suppressor.^46,47^ Beyond its enzymatic activity, RNase T2 can trigger immune responses within the tumor microenvironment, particularly by promoting the recruitment of M1-polarized macrophages.^48,49^

Notably, all HDAC6-KO models (K562, KCL22 and MV4-11) demonstrated a consistent and significant (*p* < 0.05) upregulation of RNase T2 (Figure 3A-C). Consistent with the MS-based secretome analysis, HDAC6-KO K562 cells exhibited elevated levels of RNase T2 in the culture supernatant (Figure S3A). Next, to determine whether this effect was specific to myeloid leukemia cells or extended to B-ALL cells, we assessed RNase T2 protein levels in HDAC6-KO or knockdown (KD) models using B-ALL cell lines (NALM6 and NALM20) and in (*BCR::ABL1^+^*) PDX-ALL2 cells (Figure S3B-E and Table S2). Specifically, RNAse T2 expression remained unchanged or even downregulated upon HDAC6 loss in the tested B-ALL models. Moreover, correlation analysis (using the St. Jude PeCan database) revealed that in AML patients, HDAC6 and RNase T2 expression levels are negatively correlated (r = -0.13, p < 0.05), whereas in lymphoid leukemia, a strong positive correlation (r = 0.53, p < 0.0001) was observed (Figure S3F-I).

**Figure 3.**
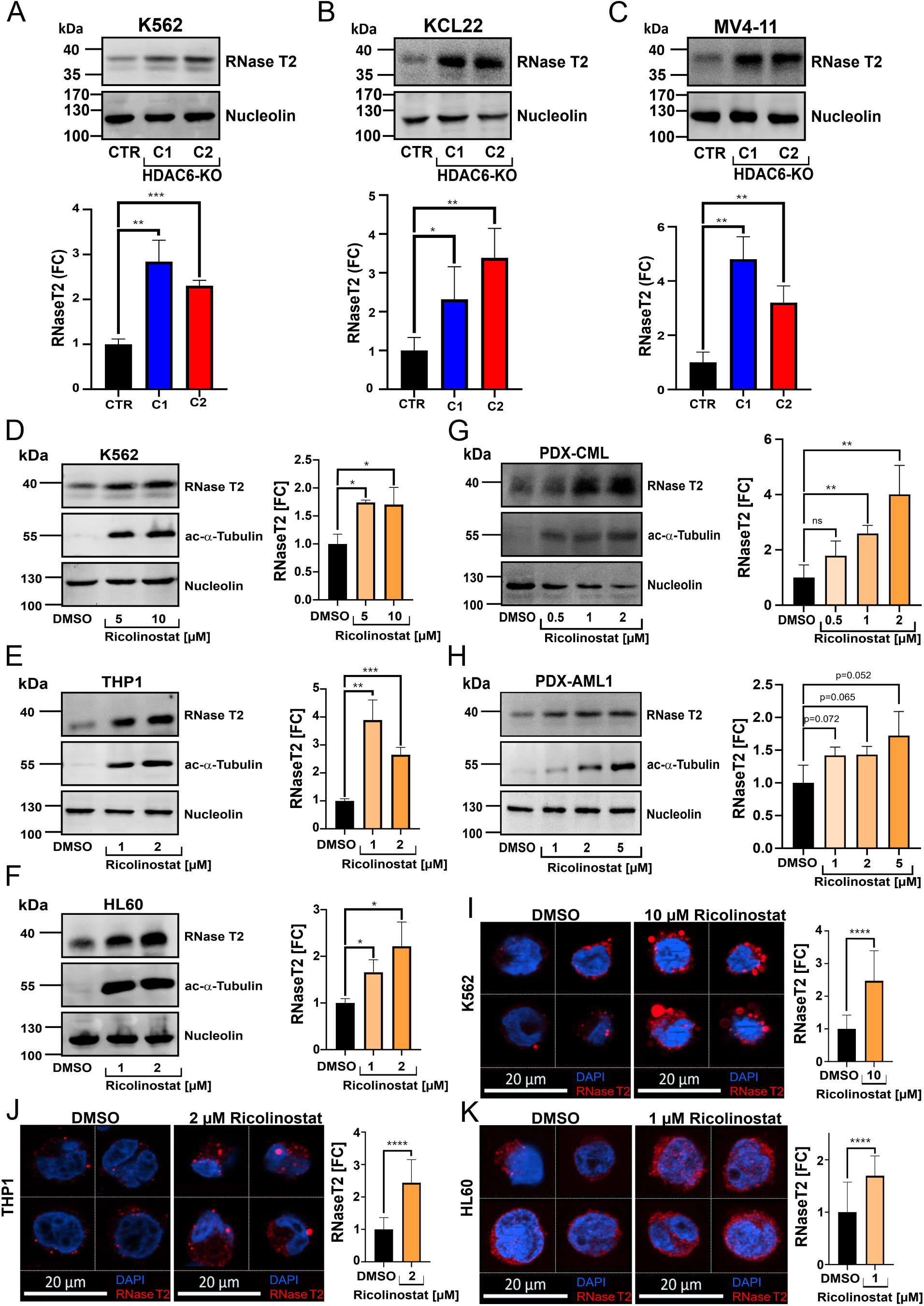
Targeting HDAC6 induces the expression of tumor suppressor RNase T2 in myeloid leukemia cells. (A-C) Western blot analysis of RNase T2 expression in HDAC6-KO myeloid leukemia models: K562 (A), KCL22 (B) and MV4-11 (C), and in the lower panel quantification results in form of a bar graph (unpaired t-test, **p<0.01, ***p<0.001, n=3). (D-H) Western blot analysis of RNase T2 and acetylated α-tubulin levels in K562 (D), THP-1 (E), HL60 (F), PDX-CML (G) and PDX-AML1 (H) cells following 24-hour treatment with the HDAC6 inhibitor Ricolinostat at the indicated concentrations. The right panel shows the results of the quantification normalized to housekeeper Nucleolin (unpaired t-test, *p<0.05, **p<0.01, ***p<0.001, n=3). (I-K) Fluorescence microscopy analysis of RNase T2 localization and expression in K562 (I), THP-1 (J), and HL60 (K) cells treated with Ricolinostat for 24 hours at the indicated concentrations. Representative cells are shown, while in the right panel the bar graph compares the fold change of fluorescence signal across all analyzed cells (unpaired t-test, ns = not significant, ****p<0.0001, n=20).

Short-term treatment with the HDAC6 inhibitor Ricolinostat at sub-cytotoxic concentrations (confirmed by an increase in acetylated tubulin levels, indicating effective HDAC6 inhibition) also resulted in a significant (p < 0.05) upregulation of RNase T2 expression across multiple myeloid leukemia cell lines, including K562, THP1, and HL60 (Figure 3D-F). Additionally, we included (*BCR::ABL1*^+^) PDX-CML and (*RUNX1-*mut) PDX-AML1 cells in the assay, which also revealed a concentration-dependent increase in RNase T2 levels upon treatment with Ricolinostat (Figure 3G-H and Table S2). Immunofluorescence microscopy further confirmed that RNase T2 accumulated in a punctate pattern upon Ricolinostat treatment, in myeloid leukemia (K562, MV4-11, THP-1, and HL60) cells (Figure 3I-K and Figure S3J). These findings suggest that HDAC6 inhibition effectively induces a phenotype characterized by a low HDAC6/RNase T2 ratio. Notably, a survival analysis (using TARGET dataset) showed that AML patients with a low HDAC6/RNase T2 ratio had significantly (*p* = 0.0003) improved survival outcomes in comparison to those with a high ratio, whereas the opposite trend was observed in ALL dataset. (Figure S3K-L). Additionally, better survival in AML was associated with lower levels of anti-inflammatory monocytes and M2 macrophages (Figure S3M-N), aligning with previous findings (in solid cancers) where HDAC6 inhibition or RNase T2 upregulation promotes macrophage polarization toward a pro-inflammatory phenotype.^47,50^ Taken together, HDAC6 regulates RNase T2 expression in myeloid leukemia cells.

### HDAC6 inhibition enhances the accessibility at the LAMP1 and RNase T2 gene loci

Although HDAC6 primarily resides in the cytoplasm, it can localize to the nucleus and participate in chromatin regulation.^51^ To investigate the mechanism underlying RNase T2 upregulation following HDAC6 loss, we performed ATAC-seq analysis. While only a limited number of genomic loci exhibited chromatin compaction following HDAC6 loss, a substantial number demonstrated increased accessibility, including loci encoding LAMP1 and RNase T2 (Figure 4A-E and Figure S4A). Next, to assess how increased chromatin accessibility influences global gene expression, we performed RNA-seq on HDAC6-KO cells (Figure S4B). Gene set enrichment analysis (GSEA) on the RNA-seq data identified upregulation of pathways related to protein acetylation regulation, exocytotic vesicle secretion, and MAPK signaling, while processes associated with HDAC6 function, including aggresome and autophagosome formation, were downregulated (Figure S4C-D).^52,53^ To integrate our findings, RNA-seq data were overlaid onto the ATAC-seq dataset, revealing a substantial (24.2%) overlap between differentially accessible chromatin regions and transcriptionally upregulated genes (Figure S4E-F). Integrated multi-omics analysis, incorporating ATAC-seq, RNA-seq, proteomic and secretomic data, revealed significant enrichment of gene sets associated with the cell periphery and plasma membrane (Figure 4F and Figure S4G). These findings suggest that HDAC6 inhibition may regulate surface receptor expression and influence leukemia cell interactions with the surrounding microenvironment.^54^

**Figure 4.**
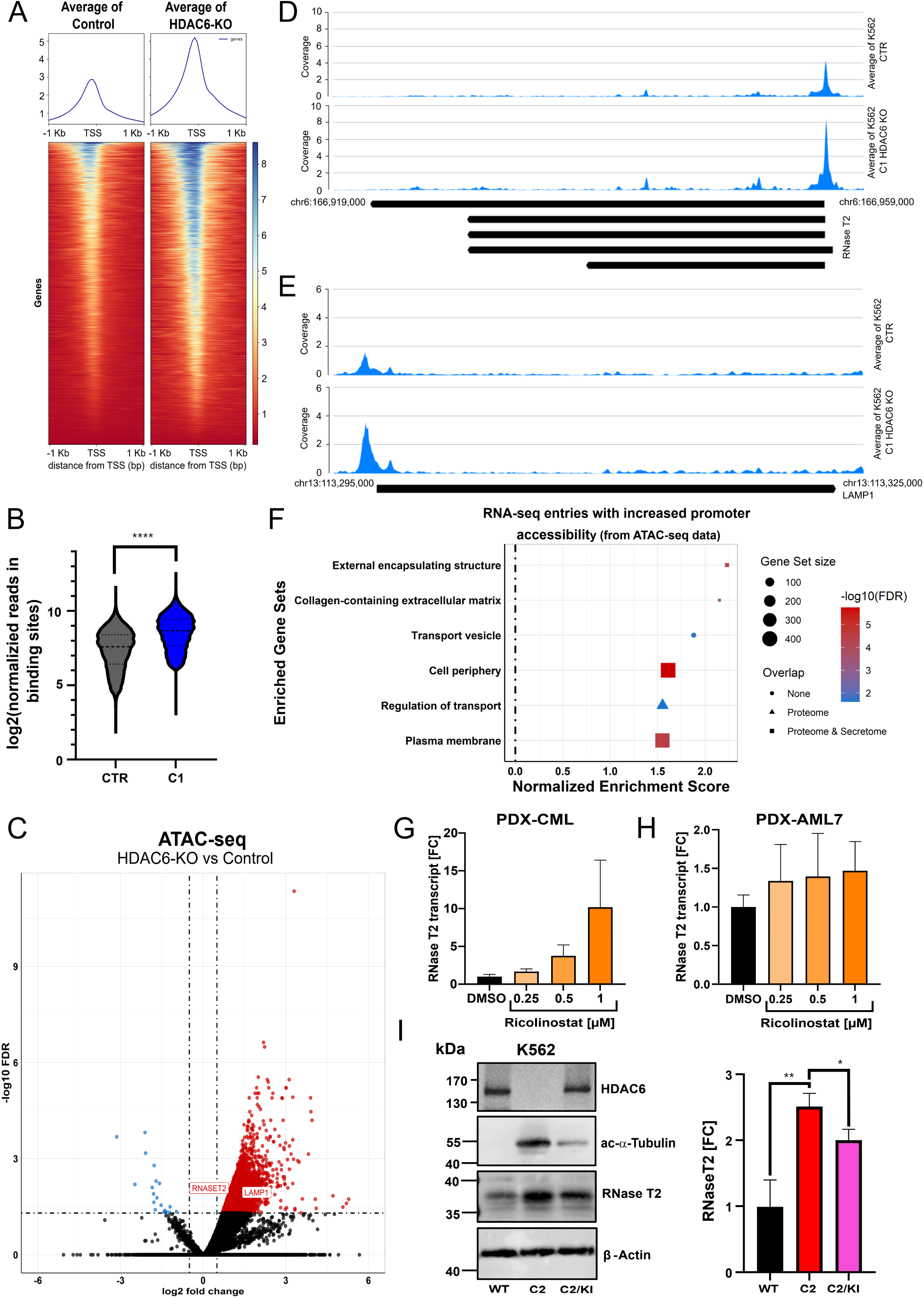
HDAC6 inhibition enhances the accessibility at the LAMP1 and RNase T2 gene loci. (A) Heatmap illustrating the distribution of chromatin accessibility relative to the transcription start site (TSS) in control (CTR) and HDAC6-KO (C1) K562 cells, as determined by ATAC-seq (n=2). (B) Violin plot comparing the log_2_-normalized read density at the binding sites of all significantly enriched genes between CTR and HDAC6-KO (Mann-Whitney U-test, ****p<0.0001, n=10,577). (C) Volcano plot depicting differentially accessible chromatin regions in HDAC6-KO C1 versus CTR cells, with the log₂ fold change plotted against the –log_10_(p-value). The cut-off thresholds were set at log₂ fold change > 0.5 and –log_10_(FDR) > 1.301 (quasi-likelihood F-test, Benjamini-Hochberg correction, n=2). (D-E) ATAC-seq track plots illustrating chromatin accessibility at the RNase T2 (D) and LAMP1 (E) loci in CTR and HDAC6-KO K562 cells. (F) Dot plot showing the manually curated enriched gene sets obtained by a *clusterprofiler* analysis for the significant RNA-seq entries with significantly increased accessibility in the promotor region (FDR<0.05). The shape of the dot indicates whether the term was also significantly upregulated in the Proteome or Secretome (Kolmogorov-Smirnov test, Benjamini-Hochberg correction). (G-H) Quantitative PCR analysis of RNase T2 expression in PDX-CML (G), and PDX-AML7 (H) samples treated with Ricolinostat for 72 hours at the indicated concentrations. Expression levels were normalized to β-actin and B2M (n=3). (I) Western blot analysis of RNase T2 acetylated α-tubulin levels in K562 control (CTR), HDAC6-KO (C2), and HDAC6-KO (C2) with HDAC6 knock-in (C2/KI). The right panel (bar graph) shows the results of the quantification normalized to housekeeper β-actin (unpaired t-test, *p<0.05, **p<0.01, n=3).

Treatment with HDAC6 inhibitors (Ricolinostat or Citarinostat) also resulted in increased RNase T2 transcription in AML-derived (*KMT2A-r*) THP1 cells, as well as in (*BCR::ABL1^+^*) PDX-CML and (*KMT2A-r*) PDX-AML7 cells, mirroring the effects observed following genetic ablation of HDAC6 (Figure 4G and Figure S4H-J). Next, to determine whether RNase T2 upregulation upon HDAC6 inhibition is reversible, we performed knock-in (KI) experiments by reintroducing HDAC6 into HDAC6-KO cells. Functional validation of HDAC6 re-expression was confirmed by the reversal of α-tubulin hyperacetylation (Figure 4I). HDAC6 restoration in HDAC6-KO cells also led to a significant (*p* < 0.05) reduction in RNase T2 expression, supporting the role of HDAC6 in the regulating RNase T2 expression. These results show that HDAC6 inhibition induces chromatin relaxation and upregulates RNase T2, an effect reversible upon HDAC6 reintroduction.

### HDAC6 inhibition sensitizes myeloid leukemia cells to CD8^+^ T cells

Based on our multi-omics and in vivo data suggesting that targeting HDAC6 affects immune-related signaling, we aimed to further explore its immunomodulatory role in a more immunocompetent in vivo setting. We therefore employed a syngeneic mouse model using C1498 cells, derived from C57BL/6 mice and classified as acute myelomonocytic leukemia.^55^ Briefly, murine C1498-Luc-GFP+ cells were injected into immunocompetent C57BL/6 wild-type mice (Figure 5A). After confirming engraftment, the mice were treated with four doses of Ricolinostat (50 mg/kg). Bone marrow, spleen and lymph nodes were collected after the last treatment. A significant increase (p < 0.05) in TNFα production was observed in bone marrow-infiltrating T cells from Ricolinostat-treated mice, whereas only a modest (non-significant) elevation was detected in splenic T cells, and no changes were observed in lymph node T cells (Figure 5B and Figure S5A– B). Similarly, CD107a (LAMP1) expression was upregulated in bone marrow T cells but remained unchanged in those from the spleen and lymph nodes (Figure 5C and Figure S5C-D), a marker associated with cytotoxic T lymphocyte (CTL) lytic granule release and target cell cytotoxicity.^56^

**Figure 5.**
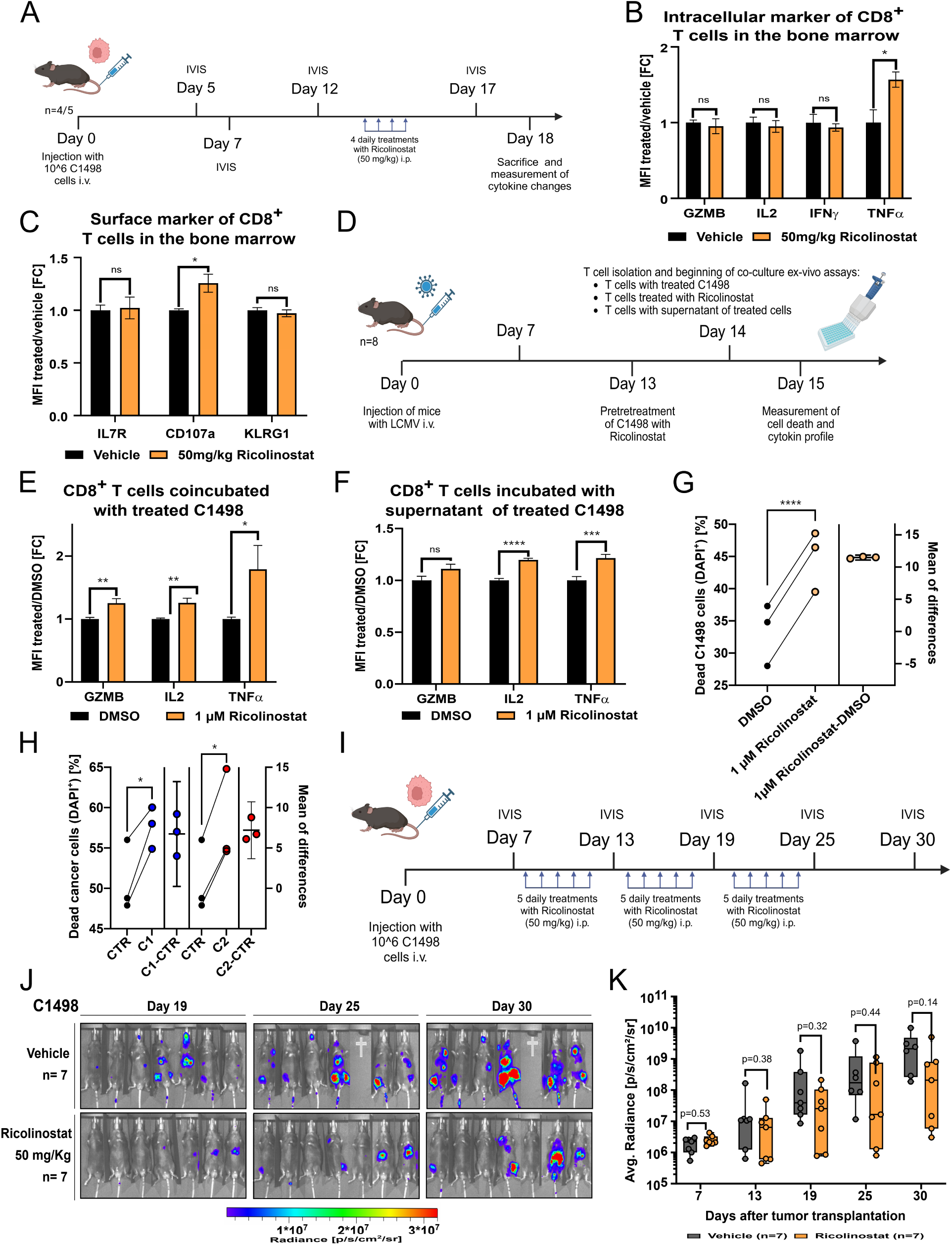
HDAC6 inhibition sensitizes myeloid leukemia cells to CD8+ T cells. (A) Experimental timeline of Ricolinostat (50 mg/kg) treatment following the injection of murine C1498-Luc-GFP+ AML cells into wild-type C57BL/6 mice to determine CD8+ T cell marker level. (B-C) Bar graph comparing the fold change in mean fluorescence intensity (MFI) of intracellular (B) or surface (C) marker of CD8^+^ T cells from the bone marrow after Ricolinostat treatment against the control (unpaired t-test, ns = not significant, *p < 0.05, n = 4-5). (D) Experimental timeline for the harvest of primed CD8^+^ T cells from C57BL/6 mice following Lymphocytic Choriomeningitis Virus (LCMV) injection, followed by ex vivo co-culture with C1498 cells. (E-F) Bar graph showing the fold change in MFI of quantified cytokines from CD8^+^ T cells co-incubated with C1498 cells treated for 24 hours with 1 µM Ricolinostat (E) or the supernatant from Ricolinostat-treated C1498 cells (F) (unpaired t-test, *p < 0.05, **p < 0.01, ***p < 0.001, ****p < 0.0001, n = 16). (G) Line plot showing the fraction of dead C1498 cells when co-incubated with CD8^+^ T cells, pretreated with Ricolinostat or DMSO (paired t-test, ****p < 0.0001, n = 3). (H) Similar to (G), but using C1498 HDAC6-KO (C1 or C2) cells vs non-targeting-control (CTR). (I) Experimental timeline of Ricolinostat (50 mg/kg) treatment following the injection of murine C1498-Luc-GFP+ AML cells into wild-type C57BL/6 mice to determine disease progression. (J) *In vivo* growth kinetics of C1498 cells in C57BL/6 mice treated with Ricolinostat, assessed at 19-, 25-, and 30-days post-injection via luminescence detection. (K) Quantification of the average radiance from (J) (Mann-Whitney U-test, n = 7).

To validate these finding, an ex vivo co-culture model was established with syngeneic murine CD8^+^ T cells (Figure 5D). Treatment of C1498 cells with Ricolinostat induced a significant (*p* < 0.01) upregulation of GZMB, IL-2 and notably TNFα in CD8^+^ T cells (Figure 5E). Moreover, the supernatant from Ricolinostat-treated C1498 cells was enough to elicit the same response in T cells, while Ricolinostat treatment alone did not significantly affect the expression of these markers (Figure 5F and Figure S5E). Notably, treatment with Ricolinostat consistently enhanced CD8^+^ T cell-mediated cytotoxicity of C1498 cells (Figure 5G). Ricolinostat treatment led to an upregulation of RNase T2 expression in murine C1498 cells, consistent with observations in human myeloid leukemia cells (Figure S5F). Additionally, immunophenotyping of Ricolinostat-treated human K562 leukemia cells revealed a concentration-dependent increase in the HLA-ABC expression (Figure S5G).

To further validate the involvement of HDAC6 in eliciting T cell responses, a C1498 HDAC6-KO model was generated (Figure S5H). While the cytokine response was less pronounced with C1498 HDAC6-KO model, we still observed a significant (*p* < 0.05) increase in CD8^+^ T cell susceptibility to cytotoxicity and cytokine responses, similar to the wild-type C1498 model upon Ricolinostat treatment (Figure 5H and Figure S5I-J). Next, to assess the therapeutic benefit of Ricolinostat as a monotherapy, we treated C57BL/6J mice following engraftment with C1498 Luc-GFP+ cells over a prolonged period (3 weeks) and monitored leukemia progression (Figure 5I). Although Ricolinostat treatment showed a trend towards restricting leukemia progression, the results were not statistically significant, indicating that while HDAC6 inhibition enhances T cell-mediated immune responses (Figure 5J-K), and its therapeutic benefit in vivo requires additional combination partners.

### Targeting HDAC6 sensitizes myeloid leukemia cells towards standard chemotherapeutics Cytarabine and Clofarabine

To enhance the therapeutic impact of HDAC6 inhibition and identify potential anti-cancer combination partners, we conducted a synthetic lethality screen using HDAC6-KO cells (Figure 6A). HDAC6-KO cells showed significantly (p < 0.05) increased sensitivity to several drug classes, including tyrosine kinase inhibitors (TKIs) and anti-metabolites such as Clofarabine, Cytarabine and 5-Azacytidine. Given the clinical relevance of Cytarabine in AML, we examined its effects alongside Clofarabine. Both drugs exhibited enhanced cytotoxicity in HDAC6-KO cells, with Clofarabine showing a particularly strong reduction in proliferation (Figure 6B and Figure S6A). HDAC6-KO cells also displayed increased apoptosis induction (p < 0.01) and sub-G1 accumulation following treatment of Cytarabine or Clofarabine (Figure 6C-D and Figure S6B-C). Elevated levels of PARP cleavage (p < 0.05) following treatment with Cytarabine or Clofarabine further confirmed enhanced apoptotic responses in HDAC6-KO cells compared to corresponding controls (Figure 6E). These findings suggest that HDAC6 loss can enhance the sensitivity to nucleoside analogs.

**Figure 6.**
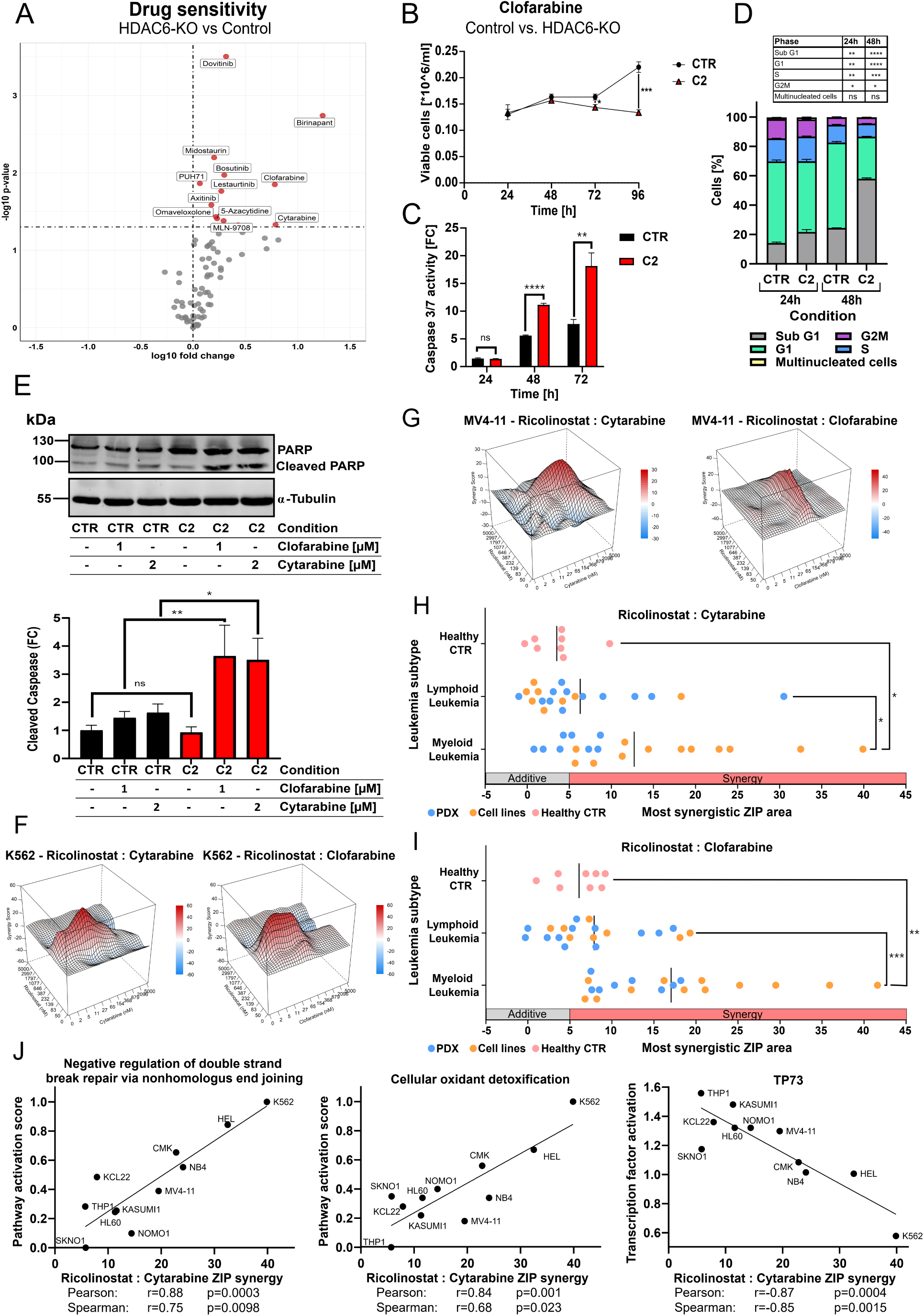
Targeting HDAC6 sensitizes myeloid leukemia cells towards standard chemotherapeutics Cytarabine and Clofarabine. (A) Volcano plot depicting the log_10_(IC50 FC) and the –log_10_(p-value) between K562 CTR and K562 HDAC6 KO (C1 and C2). As cut off was -log_10_(p-value) >1.301 (equals <0.05) used (unpaired t-test, n=3-6). (B) Proliferation curve determined by trypan blue assay of K562 CTR and K562 HDAC6-KO C2 treated with 1 µM Clofarabine over the course of 96 h (unpaired t-test, *p< 0.05, ***p< 0.001, n=3). (C) Bar plot of the fold change of the caspase 3/7 activity of K562 CTR and K562 HDAC6 KO C2 treated with 1 µM Clofarabine over the course of 72h (unpaired t-test, ns= not significant, **p < 0.05, ****p < 0.0001, n = 3). (D) Bar plot showing a cell cycle analysis of K562 HDAC6 KO models treated with Clofarabine. Statistical significance was determined via an unpaired t-test (ns= not significant, *p< 0.05, **p< 0.01, ***p< 0.001, ****p< 0.0001, n=3). (E) Western Blot analysis of PARP and cleaved PARP of K562 HDAC6 KO models treated with 1µM Clofarabine or 2 µM Cytarabine. The lower panel bar graph shows the results of the quantification of cleaved PARP normalized to the housekeeper α-tubulin (unpaired t-test, *p< 0.05, **p< 0.01, n=3). (F-G) Matrix synergy plots of K562 (F) and MV4-11 (H) cell lines depicting the combination of increasing concentrations of Ricolinostat with increasing concentrations of Cytarabine or Clofarabine. (H-I) Dot plot summarizing the most synergistic ZIP scores of AML/CML cell lines (n=12), patient-derived PDX-AML/CML cells (n=8), ALL cell lines (n=8), PDX-ALLs (n=12), and healthy controls (n=8), when treated with Ricolinostat and Cytarabine (H) or Ricolinostat and Clofarabine (I) combination. Statistical significance was determined using an unpaired t-test (ns = not significant, *p < 0.05, **p < 0.01, ***p < 0.001). (J) Pathway and transcription factor activities were inferred from gene expression data of AML and CML cell lines and correlated with ZIP synergy scores of the Ricolinostat and Cytarabine combination.

To validate these finding, we investigated potential synergistic interactions between Ricolinostat and Cytarabine or Clofarabine (Figure 6F-G). Both Cytarabine and Clofarabine exhibited strong synergistic effects with Ricolinostat in K562 and MV4-11 cells. Encouraged by these findings, we extended our investigation to a broader cohort, including (n=10) AML, (n=4) B-ALL, and (n=4) T-ALL cell lines. Additionally, we included PDX-AML/CML cells derived from initial or relapsed pediatric/adult patients (n = 8), along with PDX-B-ALL (n = 5), PDX-T-ALL (n = 7), and healthy donor-derived PBMCs (n = 4), fibroblasts (n=2), mesenchymal stem cells (MSCs), cord blood-derived CD34⁺ cells as controls (Figure 6H-I and Figure S6D-E). Notably, myeloid lineage-derived leukemia cells from demonstrated heightened sensitivity to the HDAC6 inhibitor Ricolinostat, when combined with Clofarabine or Cytarabine, compared to B-ALL, T-ALL and healthy control samples. Next, to investigate the underlying mechanisms of the observed synergy between the HDAC6 inhibitor Ricolinostat and Clofarabine or Cytarabine, we correlated publicly available baseline transcriptomic profiles (from DepMap portal, 25Q2) of AML and CML cell lines with experimentally determined ZIP synergy scores (Figure 6J and Figure S6F). This analysis revealed a positive correlation between synergistic response and gene sets involved in the negative regulation of double-strand break repair via non-homologous end joining, as well as cellular oxidant detoxification pathways. These findings suggest that impaired DNA repair capacity and elevated oxidative stress may enhance the efficacy of combination therapy. Conversely, a negative correlation was observed with TP73 expression, a member of the p53 family involved in DNA damage response and apoptosis regulation, indicating that lower TP73 levels may sensitize cells to the combined treatment. Together, these data point to a potential mechanistic link between HDAC6 inhibition, defective DNA repair, and redox vulnerability that may be exploited to enhance chemotherapeutic efficacy of HDAC6 inhibition in myeloid leukemias.^57,58^ However, additional preclinical evaluation is warranted to assess the therapeutic potential and clinical applicability of this combination strategy.

## Discussion

Despite substantial progress in elucidating the molecular pathogenesis of AML, therapeutic strategies have remained largely unchanged, relying predominantly on conventional chemotherapy and allogeneic hematopoietic stem cell transplantation.^3,59–61^ Therefore, there is a pressing need to identify and develop novel therapeutic strategies. HDAC6 has emerged as a promising anti-cancer target.^15^ High HDAC6 expression correlates with poor survival in AML patients, while in vitro (*DepMap*) and pharmacologic inhibition data show limited benefit of targeting HDAC6,^28–30^ highlighting a potential disconnect between its prognostic relevance and therapeutic impact in vitro. Our in vivo findings using immunodeficient NSG mice revealed a significant reduction in leukemic progression following HDAC6 knockout, suggesting that HDAC6 loss impairs leukemogenesis, potentially via modulation of innate immune pathways that remain active in this model.^62^ In line with the above, multi-omics analyses revealed activation of several immune-related pathways following HDAC6 loss, especially RNase T2, an endoribonuclease linked to pro-inflammatory macrophage activation within the tumor microenvironment.^48,49^ Extending our findings to immunocompetent models, we found that HDAC6 inhibition enhanced CD8^+^ T cell activation,^63^ through TNFα induction and upregulation of CD107a (or LAMP1). Consistently, we have observed an increase in lysosome-associated proteins upon targeting HDAC6, which aligns with emerging evidence that the lysosome plays a crucial role in regulating the cytotoxic activity of CD8+ T cell immunity.^64^ Additionally, HDAC6 loss sensitized AML cells to nucleoside analogs Cytarabine and Clofarabine. This is consistent with previous findings demonstrating that dual inhibition of HDAC1 and HDAC6 potentiates the response of AML cell lines to Cytarabine.^45^ These effects were replicated with pharmacological inhibition of HDAC6 and supported by our proteomics data showing downregulation of DNA repair pathways in HDAC6-KO cells. ^57,58^ Owing to the heterogeneous expression of HDAC6 across AML subtypes, future studies should aim to identify molecularly defined patient subgroups with increased sensitivity to HDAC6 inhibition. Our data mining analyses revealed that high HDAC6 expression is particularly associated with poor prognosis in AML cases harboring NRAS mutations, as well as in FLT3- and IDH1-mutated subtypes (Fig S1E-H), warranting further investigation into these specific genetic contexts.

Together, our study provides compelling evidence that HDAC6 inhibition enhances immune-mediated leukemia cell clearance and sensitizes myeloid leukemia cells to chemotherapeutic agents. Given the minimal toxicity associated with HDAC6 inhibition, these findings support its potential use in combination with emerging immunotherapeutic strategies to enhance anti-leukemic efficacy.

## Supporting information

Supplement differential gene analysis

Supplement gene set enrichment & string analysis

## Acknowledgements

A.B. acknowledges the financial support from Katharina-Hardt Foundation, Christiane und Claudia Hempel foundation and Löwenstern e.V.

## Author Contributions

J.S-D., J.T. and S.B. performed study concept and design. J.S-D., J.T., P.S., K.S., M.K. E.V., N.R., M.V., D.B., T.L., A.N. and S.B. performed development of methodology, investigation, analysis and interpretation of data. S.S., P.D., A.S., K.N., K.S., U.F., A.A.P and A.B. contributed to resource provision, supervision, funding acquisition, and project administration. J.S-D, J.T., A.B. and S.B. performed writing, reviewing and editing of the paper. S.B. supervised this study. All authors discussed the results and commented on the manuscript.

