## Supplement Figures for "Epigenetic remodeling via HDAC6 inhibition amplifies anti-tumoral immune responses in myeloid leukemia cells"

[1]Department of Pediatric Oncology, Hematology and Clinical Immunology, Medical Faculty, Heinrich Heine University Düsseldorf, Düsseldorf, Germany
[2]Institute of Clinical Chemistry and Clinical Pharmacology, University Hospital Bonn, 53127 Bonn, Germany
[3]Molecular Proteomics Laboratory, Biological Medical Research Center, Heinrich-Heine-University Düsseldorf, Düsseldorf, Germany
[4]Institute of Medical Microbiology and Hospital Hygiene, Heinrich Heine University Düsseldorf, Düsseldorf, Germany
[5]Institute of Immunology, Rostock University Medical Center, Rostock, Germany
[6]Université Côte d'Azur, INSERM, C3M U1065, 06204 Nice, France
[7]Department of Pediatrics, Oncology and Hematology, and Department of Genetic Predisposition to Cancer, Medical University of Lodz, Lodz, Poland
[8]St. Anna Children's Cancer Research Institute (CCRI), Vienna, Austria
[9]Labdia Labordiagnostik, Vienna, Austria
[10]Institute for Molecular Medicine, Proteome Research, University Hospital and Medical Faculty, Heinrich-Heine-University Düsseldorf, Düsseldorf, Germany
[11]German Cancer Consortium (DKTK), partner site Essen/Düsseldorf, Düsseldorf, Germany
[12]German Center for Infection Research (DZIF), Partner Site Bonn-Cologne, 53127 Bonn, German
[†] contributed equally to this work

**Running Title:**
HDAC6 inhibition immunomodulates AML cells

***Corresponding author**: Sanil Bhatia, Department of Pediatric Oncology, Hematology and Clinical Immunology, Heinrich Heine University Düsseldorf, Germany, Moorenstraße 5, Düsseldorf, 40225, Germany. Phone (+49) 211 81 04896; Fax; (+49) 211 81 16436

### Table of Contents

1. **Supplemental Figures (1-6) and Tables (1-2) .............................................. 3-18**

2. **Supplemental Materials and Methods ....................................................... 19-26**

3. **Supplemental References .......................................................................... 27-28**

### 1. Supplementary Figures:

#### Fig. S1

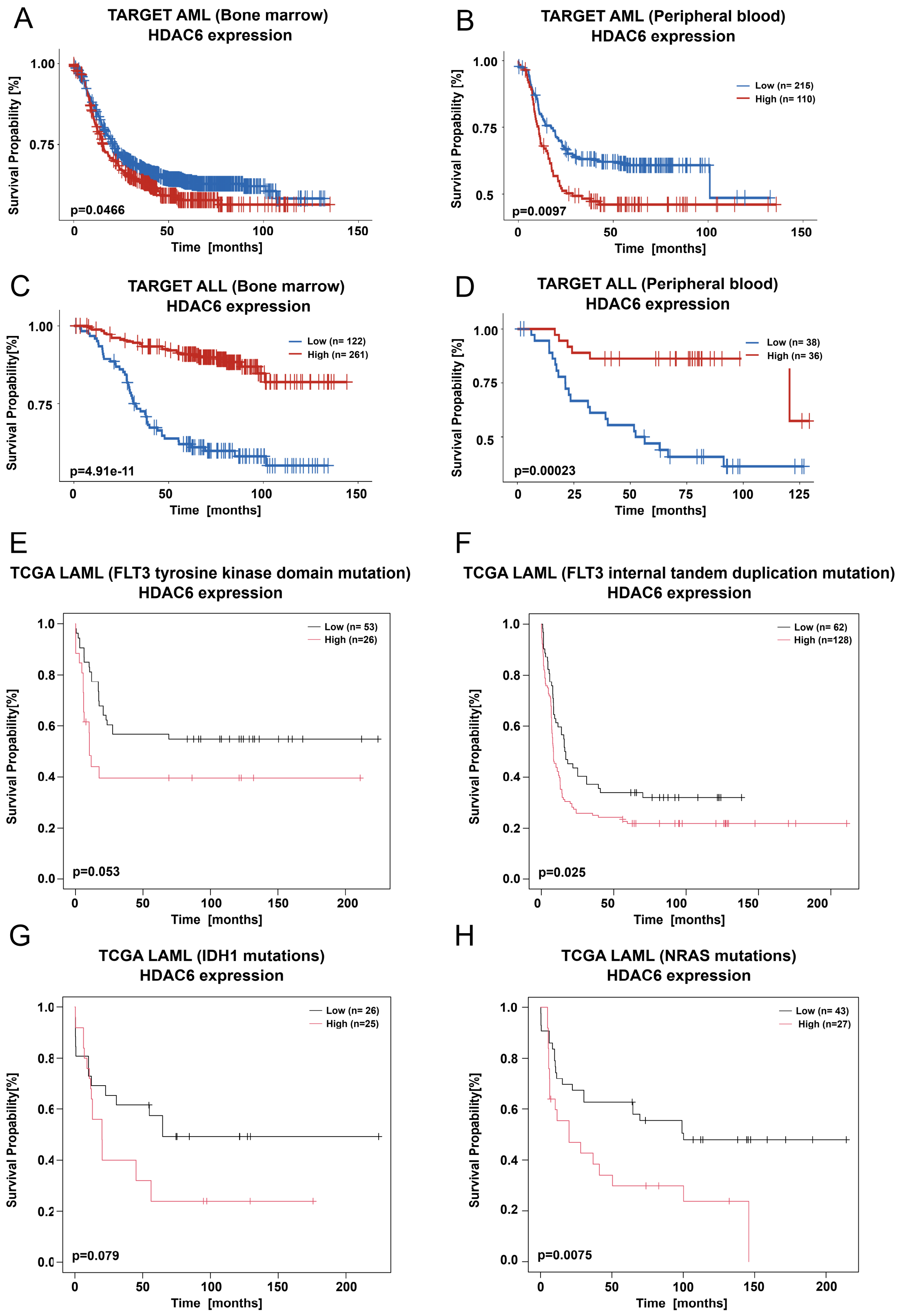

### Supplemental Figure 1B

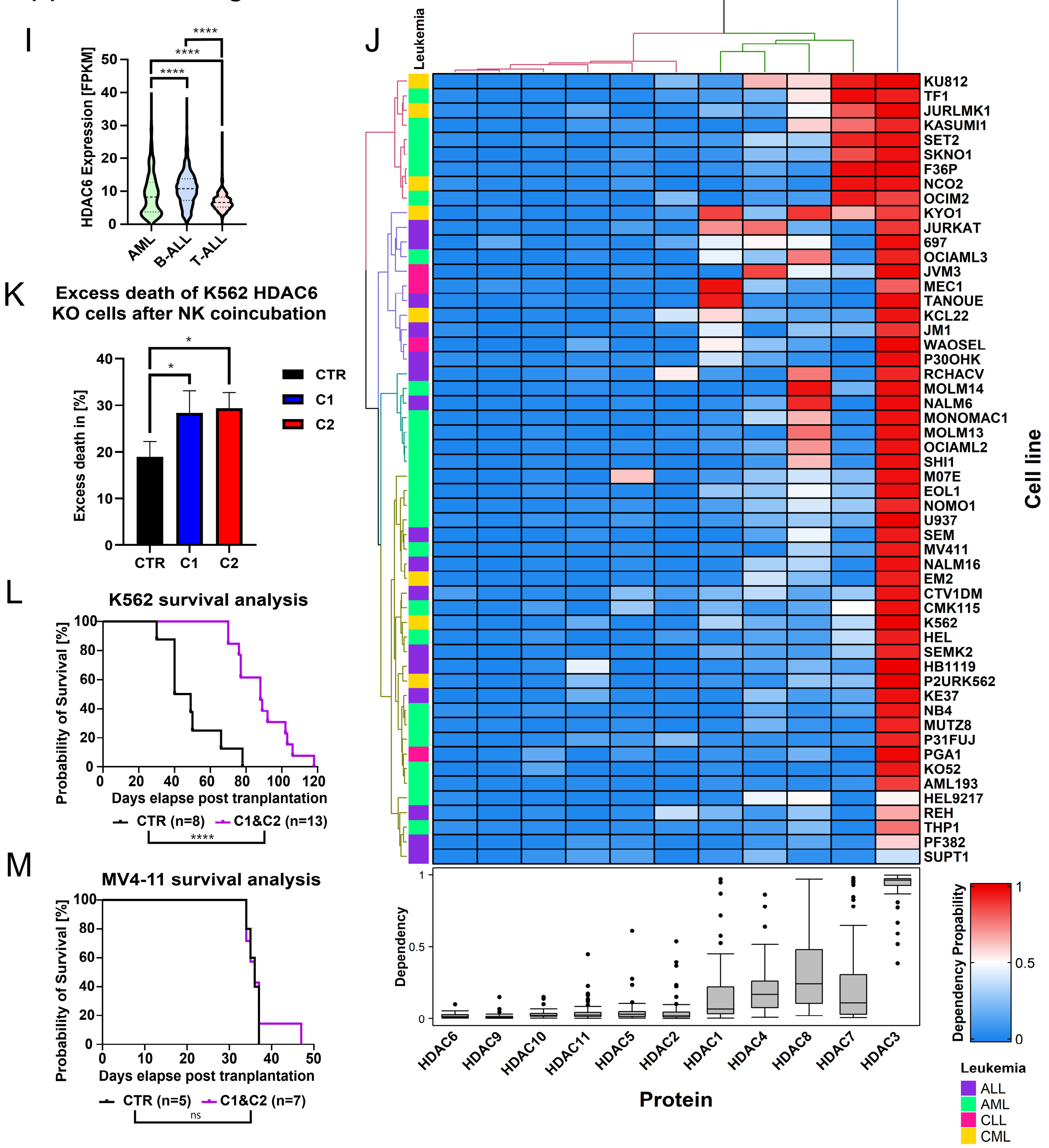


**Supplementary Figure 1. HDAC6 ablation impairs the in vivo growth of myeloid leukemia cells.** (A-D) Kaplan-Meier survival plots of AML (A-B) or ALL (C-D) patients from the *TARGET* dataset, categorized by high or low HDAC6 expression levels in bone marrow (A or C) samples or peripheral blood (B or D) (log-rank test). (E-H) Kaplan-Meier survival plots of AML categorized by high or low HDAC6 expression levels from the TCGA-LAML dataset. Patients with FLT3, IDH1 or NRAS mutation were split by a percentile-based best cut-off selection (log-rank test). (I) Violin plot displaying HDAC6 expression across the most common childhood leukemia subtypes, from the St. Jude *PeCan* database (Mann-Whitney U-test, ****p< 0.0001, n=322-813). (J) Heatmap illustrating the dependency of leukemia cell lines from different subtypes on HDAC isoforms, based on data from the DepMap portal. (K) Bar graph comparing the cell death mediated by NK cells in K562 HDAC6 KO (C1 and C2) vs the non-targeting control (CTR) (unpaired t-test, *p< 0.05, n=3). (L-M) Kaplan Meyer survival plot of mice injected with K562 (L) or MV4-11 (M) HDAC6 KO models (log-rank test, ns= not significant, ****p< 0.0001, n=5-8).

### Fig. S2

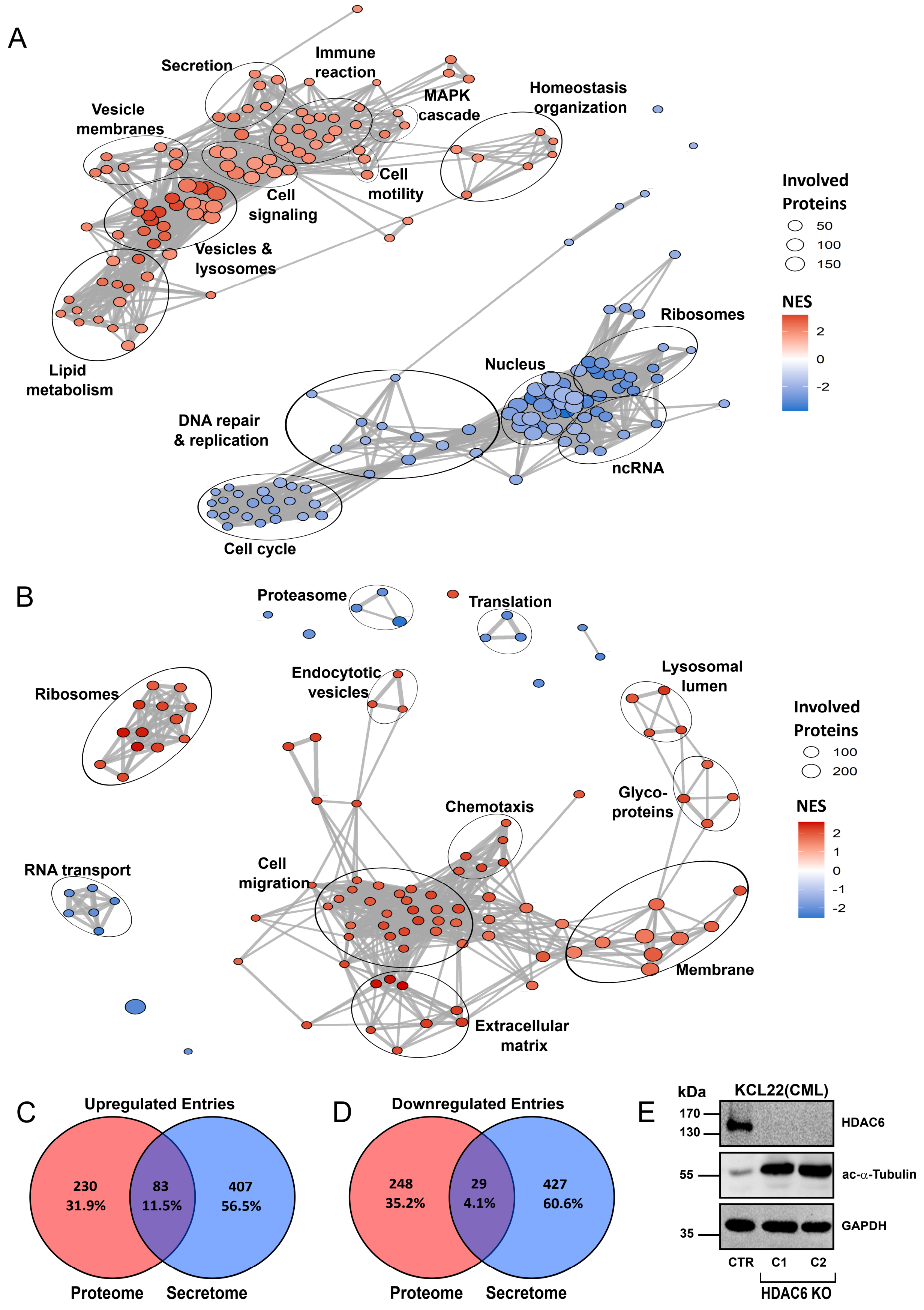
A
Secretion
Immune reaction
MAPK cascade
Homeostasis organization
Vesicle membranes
Cell signaling
Cell motility
Vesicles & lysosomes
Lipid metabolism
Involved Proteins
50
100
150
NES
2
0
-2
Ribosomes
Nucleus
DNA repair & replication
ncRNA
Cell cycle
B
Proteasome
Translation
Ribosomes
Endocytotic vesicles
Lysosomal lumen
Involved Proteins
100
200
NES
2
1
0
-1
-2
Chemotaxis
Glyco-proteins
RNA transport
Cell migration
Membrane
Extracellular matrix
C
Upregulated Entries
230
31.9%
83
11.5%
407
56.5%
Proteome
Secretome
D
Downregulated Entries
248
35.2%
29
4.1%
427
60.6%
Proteome
Secretome
E
kDa
KCL22(CML)
170
130
HDAC6
55
ac-α-Tubulin
35
GAPDH
CTR
C1
C2
HDAC6 KO

**Supplementary Figure 2. Targeting HDAC6 upregulates lysosome-related protein accumulation.** (A-B) Enrichment map of gene sets from the proteomics (A) or secretomics (B) data of K562 HDAC6 KO models, generated using *clusterProfiler*, with a protein entry inclusion criteria of FDR < 0.1. Included gene sets possessed a FDR<0.05 (Kolmogorov-Smirnov test, Benjamini-Hochberg correction). (C-D) Venn diagram showing the number of upregulated (C) and downregulated (D) overlapping and non-overlapping proteins from the proteomics and secretomics datasets with FDR < 0.1. (E) Representative western blot (WB) for HDAC6 and acetylated α-tubulin protein levels in two clones of KCL22 HDAC6 knock out (KO) (C1 & C2) and a non-targeting control (CTR, n=3).

### Fig. S3

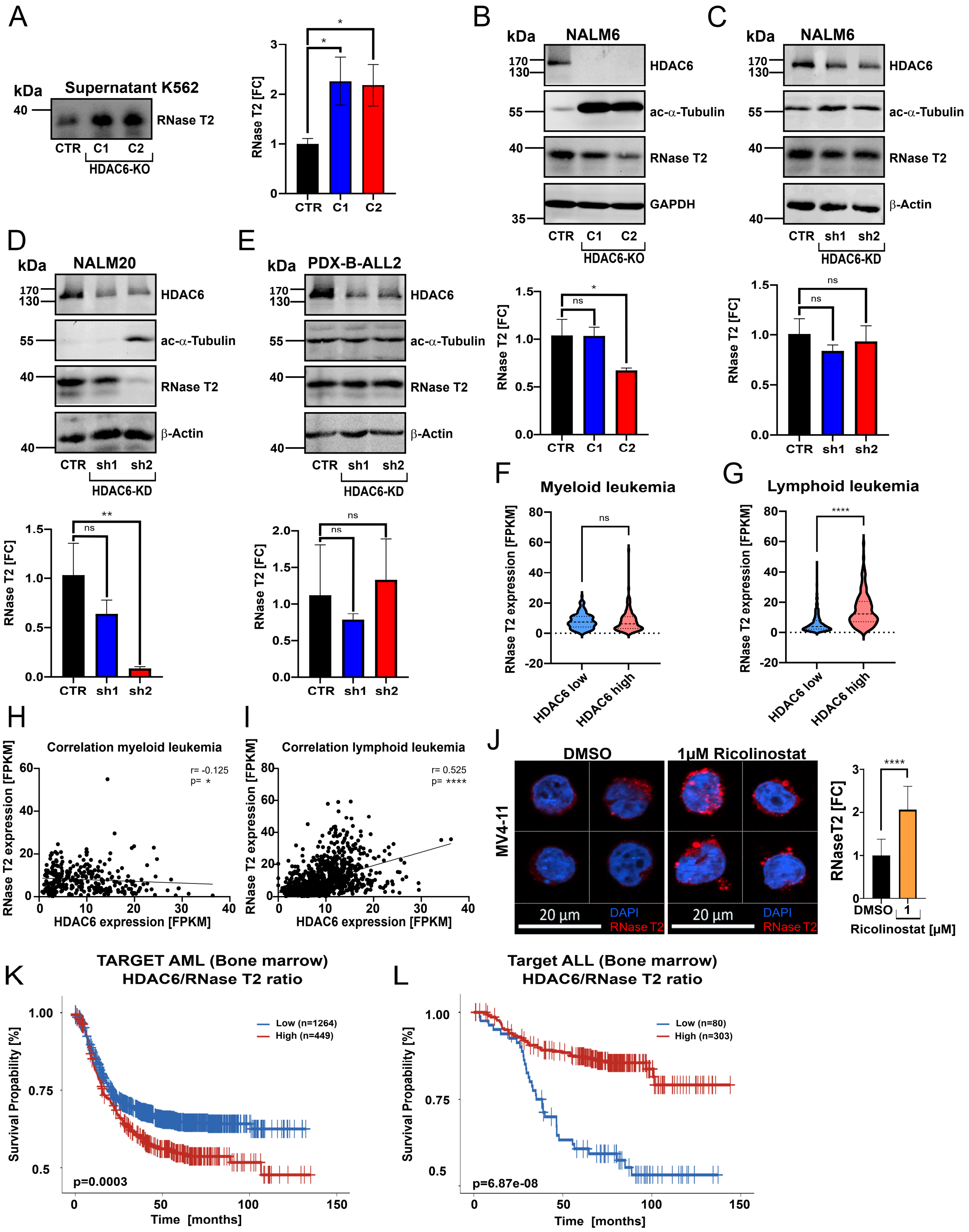
A
kDa
Supernatant K562
RNase T2
CTR C1 C2
HDAC6-KO
RNase T2 [FC]
B
NALM6
HDAC6
ac-α-Tubulin
RNase T2
GAPDH
C
β-Actin
CTR sh1 sh2
HDAC6-KD
D
NALM20
E
PDX-B-ALL2
F
Myeloid leukemia
G
Lymphoid leukemia
RNase T2 expression [FPKM]
HDAC6 low
HDAC6 high
H
Correlation myeloid leukemia
r= -0.125
p= *
HDAC6 expression [FPKM]
I
Correlation lymphoid leukemia
r= 0.525
p= ****
J
DMSO
1µM Ricolinostat
MV4-11
20 µm
DAPI
RNase T2
RNaseT2 [FC]
Ricolinostat [µM]
K
TARGET AML (Bone marrow)
HDAC6/RNase T2 ratio
Low (n=1264)
High (n=449)
p=0.0003
Survival Propability [%]
Time [months]
L
Target ALL (Bone marrow)
HDAC6/RNase T2 ratio
Low (n=80)
High (n=303)
p=6.87e-08

### Fig. S3 continued

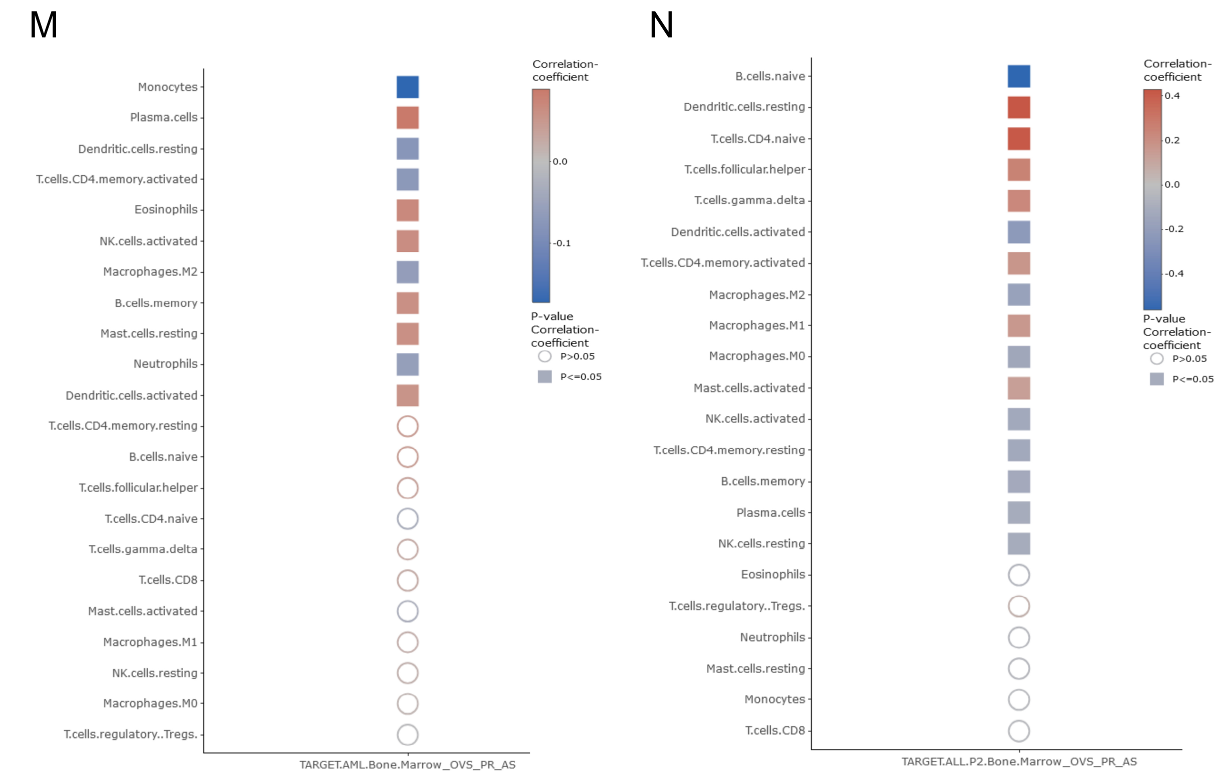


**Supplementary Figure 3. Targeting HDAC6 induces the expression of tumor suppressor RNase T2 in myeloid leukemia cells.** (A) Western blot analysis of RNase T2 levels in the supernatant of HDAC6-KO K562 cells compared to the respective control, and in right panel quantification results in form of bar graph (unpaired t-test, ns = not significant, *p < 0.05, n = 3). (B-E) Western blot analysis of HDAC6, acetylated α-tubulin, and RNase T2 levels in NALM6 HDAC6-KO (B), NALM6 HDAC6-knockdown (KD) (C), NALM20 HDAC6-KD (D) and (*BCR::ABL1*⁺) PDX-ALL2 HDAC6-KD (E) models. The lower panels shows the results of the quantification, normalized to housekeeper GAPDH or β-Actin (unpaired t-test, ns = not significant, *p < 0.05, **p < 0.01, n = 3). (F-G) Violin plots comparing the expression levels of RNase T2 in HDAC6 high and low expressing myeloid (F) or lymphoid leukemia samples (G) from the St. Jude *PeCan* database, clustered based on the mean value (Mann-Whitney U-test, ns = not significant, ****p < 0.0001, n = 160-161 for myeloid, n = 479 for lymphoid). (H-I) Dot plots showing the correlation analysis of the data from (F-G) (Spearman correlation, *p < 0.05, ****p < 0.0001, n = 321 for myeloid, n = 958 for lymphoid). (J) Fluorescence microscopy analysis of RNase T2 localization and expression in MV4-11 cells treated with Ricolinostat for 24 hours at the indicated concentrations. Representative cells are shown, while in the right panel the bar graph compares the fold change of fluorescence signal across all analysed cells (unpaired t-test, ****p<0.0001, n=20). (K-L) Kaplan-Meier survival plots of patients from the TARGET AML (K) or TARGET ALL (L) datasets, categorized by high or low HDAC6/RNase T2 expression ratio in the bone marrow (log-rank test n = 449-1264 for AML, n = 80-303 for ALL). (M-N) Correlation analysis of immune signatures from the data in (I-J) (univariate Cox proportional hazards regression model).

### Fig. S4

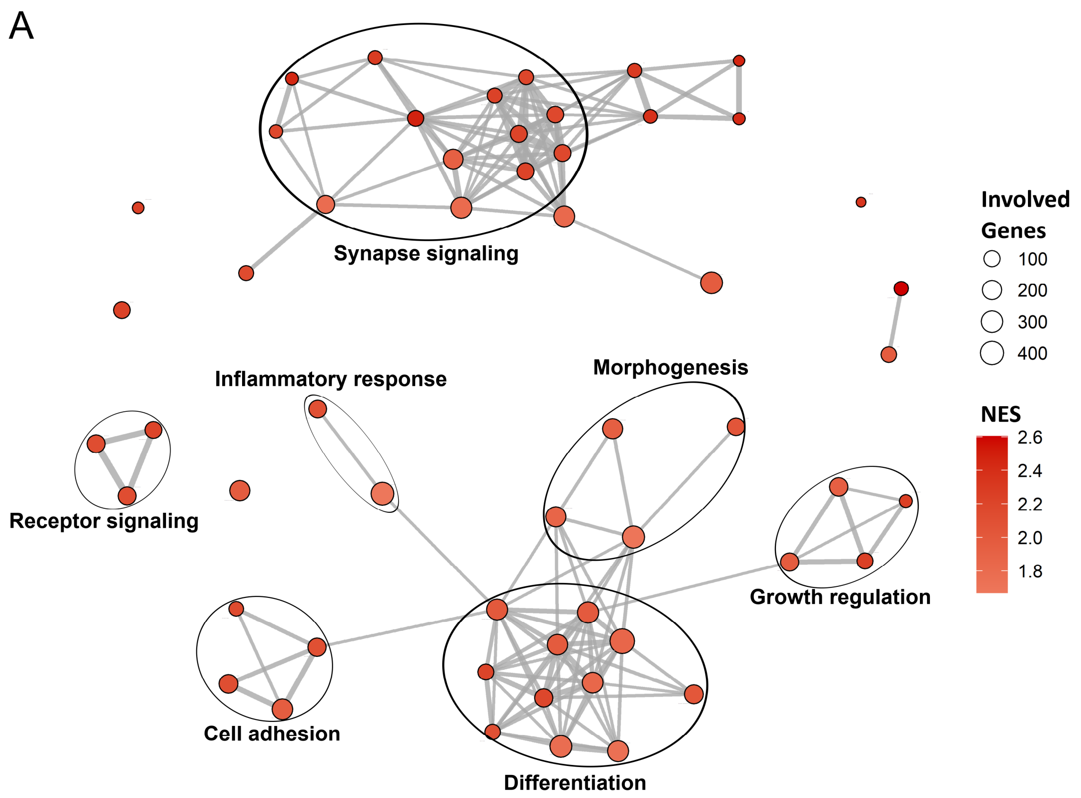
A
Synapse signaling
Inflammatory response
Morphogenesis
Receptor signaling
Growth regulation
Cell adhesion
Differentiation
Involved Genes
100
200
300
400
NES
2.6
2.4
2.2
2.0
1.8

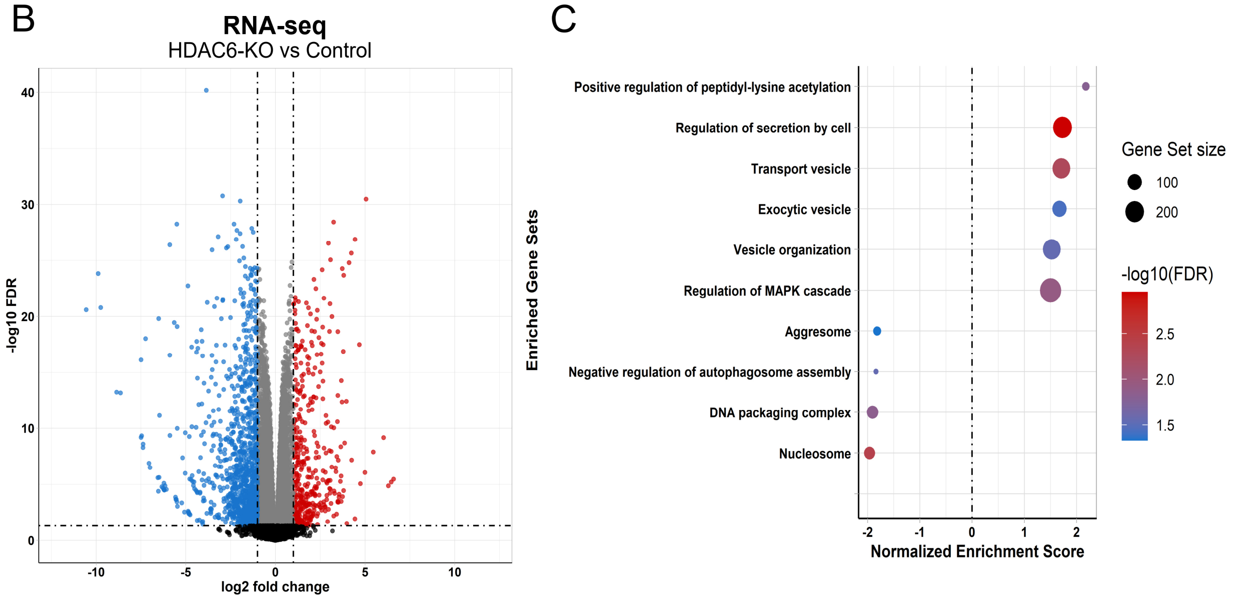
B
RNA-seq
HDAC6-KO vs Control
-log10 FDR
log2 fold change
C
Positive regulation of peptidyl-lysine acetylation
Regulation of secretion by cell
Transport vesicle
Exocytic vesicle
Vesicle organization
Regulation of MAPK cascade
Aggresome
Negative regulation of autophagosome assembly
DNA packaging complex
Nucleosome
Enriched Gene Sets
Normalized Enrichment Score
Gene Set size
100
200
-log10(FDR)
2.5
2.0
1.5

#### Fig. S4 continued

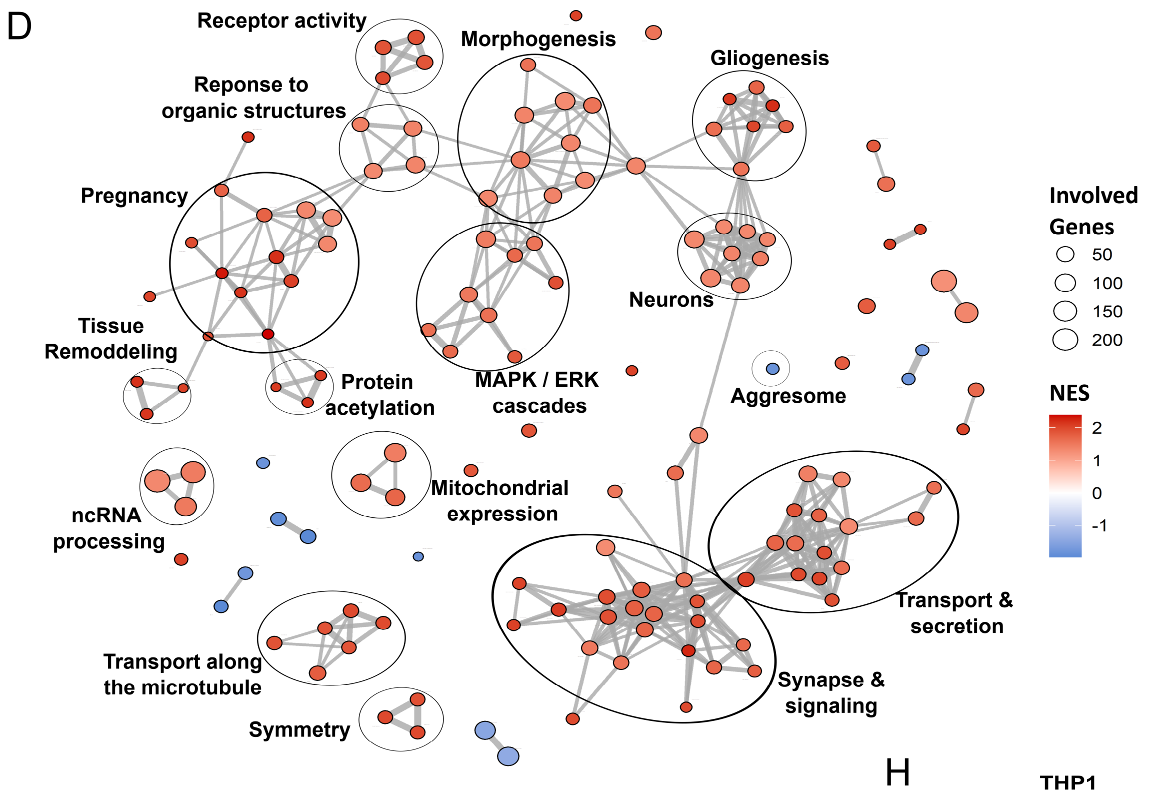


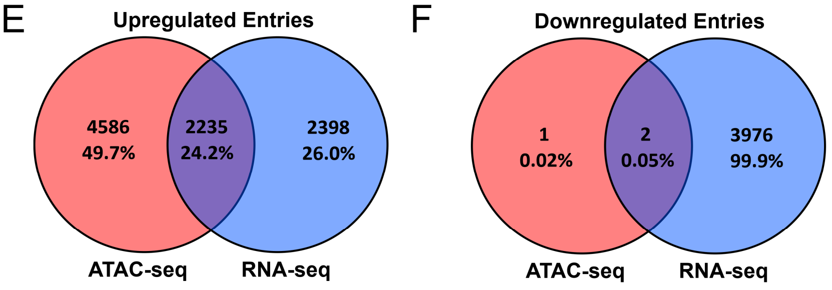


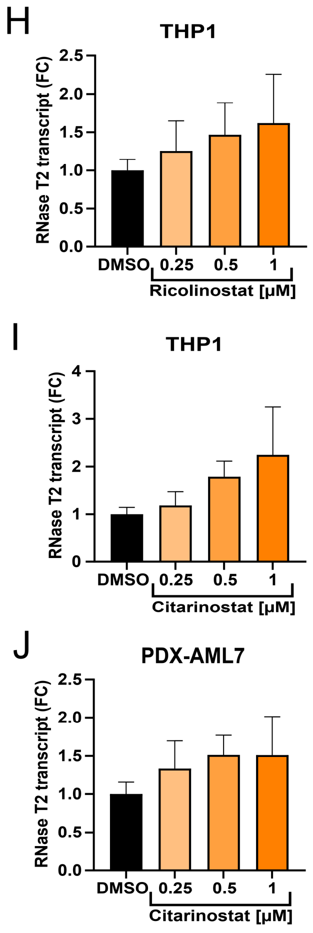


G

| Description | set Size | NES | p-value | FDR | Proteomics | Secretomics |
|---|---|---|---|---|---|---|
| Cell periphery | 403 | 1.61 | 8.51E-10 | 1.79E-06 | x | x |
| Plasma membrane | 370 | 1.55 | 4.88E-08 | 3.42E-05 | x | x |
| External encapsulating structure | 27 | 2.23 | 1.37E-07 | 5.78E-05 | x | x |
| Extracellular matrix | 27 | 2.23 | 1.37E-07 | 5.78E-05 | x | x |
| Collagen-containing extracellular matrix | 24 | 2.15 | 3.16E-06 | 5.54E-04 | x | x |
| Multicellular organismal process | 520 | 1.34 | 1.58E-04 | 1.44E-02 | x | x |
| Extracellular region | 241 | 1.41 | 1.00E-03 | 3.70E-02 | x | x |
| Cell-substrate adhesion | 31 | 1.80 | 1.38E-03 | 4.28E-02 | x | x |
| Cell communication | 459 | 1.47 | 6.68E-07 | 1.76E-04 | x | o |
| Signaling | 453 | 1.46 | 2.46E-06 | 4.71E-04 | x | o |
| Regulation of biological quality | 221 | 1.51 | 1.12E-04 | 1.21E-02 | x | o |
| Signaling receptor binding | 108 | 1.62 | 2.67E-04 | 2.16E-02 | x | o |
| Regulation of transport | 132 | 1.55 | 4.63E-04 | 2.43E-02 | x | o |
| Cell-cell signaling | 127 | 1.87 | 3.16E-08 | 3.32E-05 | o | o |
| Transport vesicle | 36 | 1.88 | 2.89E-04 | 2.17E-02 | o | o |
| G protein-coupled receptor signaling pathway | 51 | 1.73 | 4.41E-04 | 2.43E-02 | o | o |
| Cell morphogenesis | 73 | 1.61 | 1.48E-03 | 4.46E-02 | o | o |

**Supplementary Figure 4. HDAC6 inhibition enhances the accessibility at the LAMP1 and RNase T2 gene loci.** (A) Enrichment map of gene sets from the ATAC-seq data of differentially accessible promotor regions of K562 HDAC6-KO models, generated using *clusterProfiler*, with a gene entry inclusion criteria of FDR < 0.05. Included gene sets possessed a FDR<0.05 (Kolmogorov-Smirnov test, Benjamini-Hochberg correction). (B) Volcano plot depicting differentially expressed genes from RNA sequencing HDAC6-KO C1 versus K562 control cells, with the $\log_2$ fold change plotted against the –log10(p-value). The cut-off thresholds were set at $\log_2$ fold change > 1 and –log10 (FDR) > 1.301 (quasi-likelihood F-test, Benjamini-Hochberg correction, n=3). (C) Dot plot of selected enriched gene sets identified via *clusterProfiler* analysis of RNA-seq data. Significance cut-off for gene entry inclusion was set at FDR < 0.05. (Kolmogorov-Smirnov test, Benjamini-Hochberg correction). (D) Enrichment map of gene sets from the RNA-seq data, generated using *clusterProfiler*, with a gene entry inclusion criteria of FDR < 0.05. Included gene sets possessed a FDR<0.05 (Kolmogorov-Smirnov test, Benjamini-Hochberg correction). (E-F) Venn diagrams illustrating the overlap between upregulated (E) and downregulated (F) genes identified by RNA-seq and ATAC-seq analysis. (G) Table containing the curated results of a clusterprofiler GSEA of significant RNA-seq entries (FDR<0.05) with increased accessibility in the promotor region and overlap in the proteome or secretome analysis (Kolmogorov-Smirnov test, Benjamini-Hochberg correction). (H-I) Quantitative PCR analysis of RNase T2 expression in THP1 cells treated with Ricolinostat (H) or Citarinostat (I) for 72 hours at the indicated concentrations. Expression levels were normalized to β-actin and B2M (n=3). (J) Quantitative PCR analysis of RNase T2 expression in PDX-AML7 cells treated with Citarinostat for 72 hours at the indicated concentrations. Expression levels were normalized to β-actin and B2M (n=3).

### Fig. S5

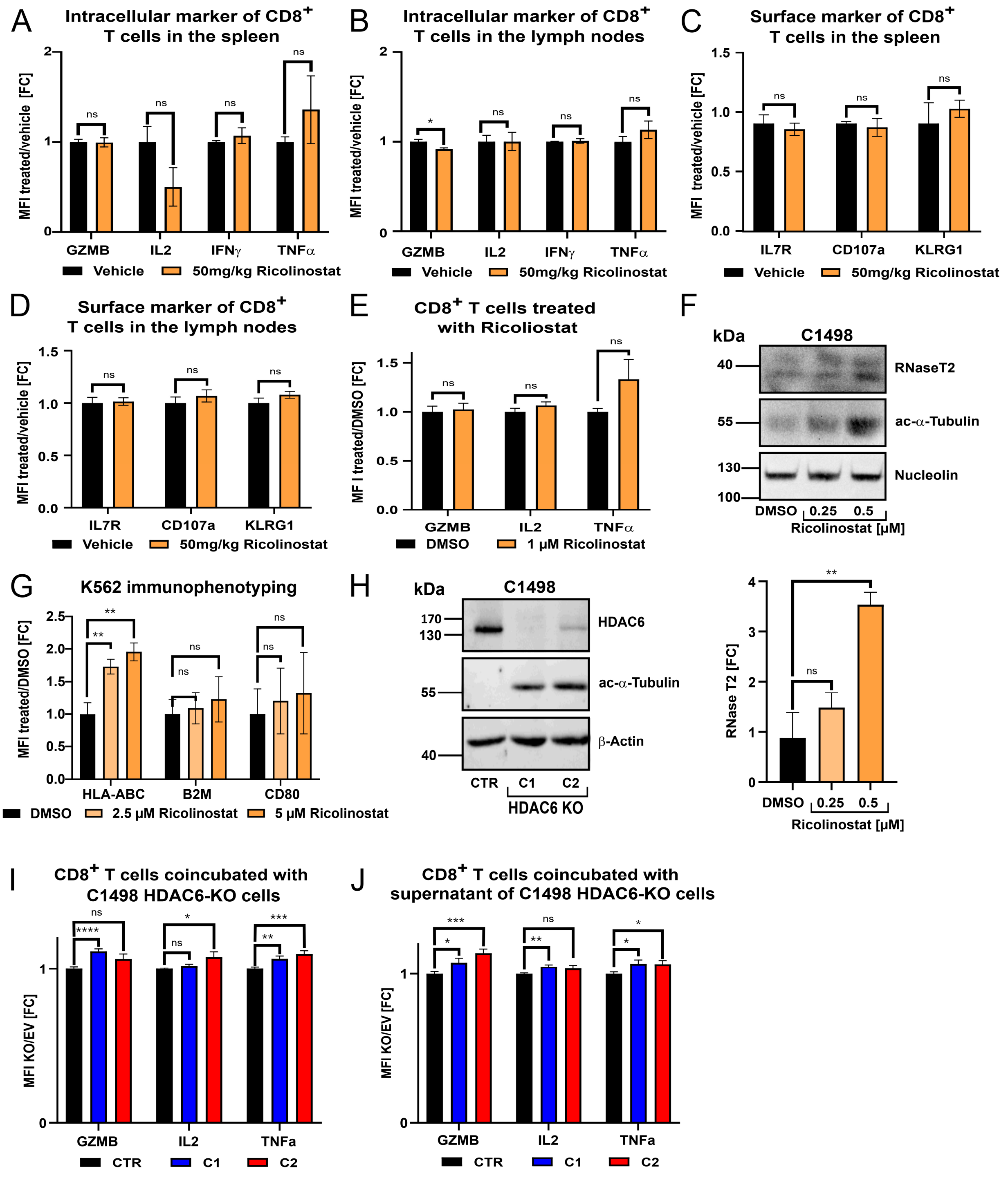


**Supplementary Figure 5. HDAC6 inhibition sensitizes myeloid leukemia cells to CD8+ T cells.** (A-B) Bar graph comparing the fold change in mean fluorescence intensity (MFI) of intracellular marker of CD8+ T cells from the spleen (A) or the lymph nodes (B) after Ricolinostat treatment against the control, following the injection of murine C1498-Luc-GFP+ AML cells into wild-type C57BL/6 mice (unpaired t-test, ns = not significant, *$p < 0.05$, n = 4-5). (C-D) Bar graph comparing the fold change in MFI of surface marker of CD8+ T cells from the spleen (C) or the lymph nodes (D) after Ricolinostat treatment against the control, following the injection of murine C1498-Luc-GFP+ AML cells into wild-type C57BL/6 mice (unpaired t-test, ns = not significant, n = 4-5). (E) Bar plot depicting fold change in MFI levels of intracellular marker of CD8+ T cells after 24 hour treatment with 1 µM Ricolinostat. (unpaired t-test, ns = not significant, n = 3). (F) Western blot analysis of RNase T2 and acetylated α-tubulin levels in C1498 cells treated for 24 hours with the indicated concentrations of Ricolinostat. The lower panel shows the results of the quantification normalized Nucleolin to as a bar graph (unpaired t-test, **p<0.01, n=3). (G) Bar graph comparing the fold change in MFI of K562 cells treated with Ricolinostat for 24 hours (unpaired t-test , ns= not significant, **p < 0.01, n = 3). (H) Representative western blot (WB) for HDAC6 and acetylated α-tubulin protein levels in two clones of C1498 HDAC6 knockout (KO) (clone C1 & C2) and a non-targeting control (CTR, n=3). (I-J) Bar graph showing the fold change in MFI of intracellular marker of CD8+ T cells co-incubated with C1498 HDAC6 KO cells (I) or their supernatant (J) (unpaired t-test, *p < 0.05, **p < 0.01, ***p < 0.001, ****p < 0.0001, n = 16).

### Fig.S6

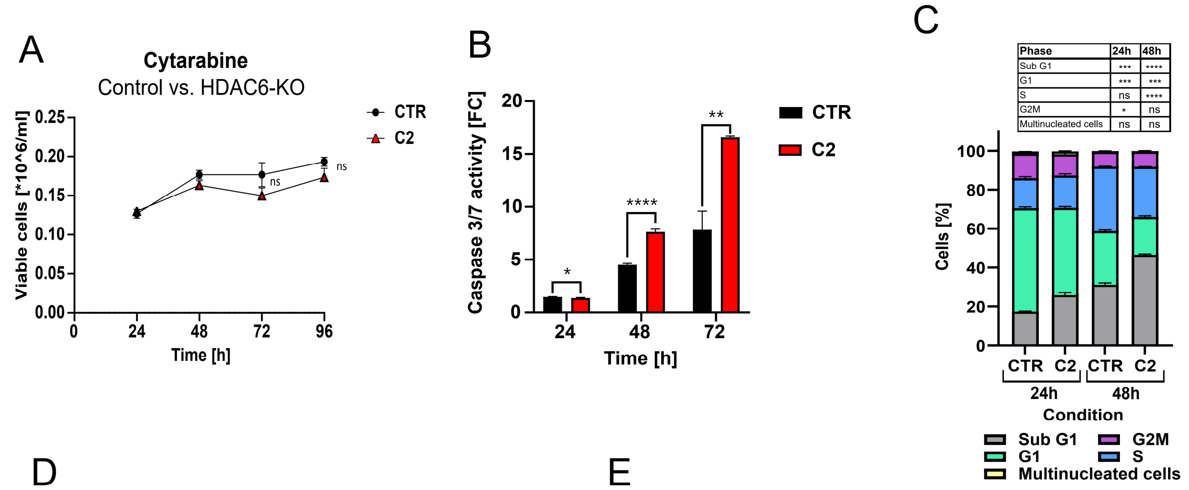


D

**Ricolinostat x Cytarabine ZIP area**

| Myeloid Leukemia | | B-ALL | | Healthy CTR | |
|---|---|---|---|---|---|
| Name | ZIP | Name | ZIP | Name | ZIP |
| HL60 | 5.7 | HAL01 | -0.1 | PBMC1 | 4.1 |
| K562 | 39.9 | MHH-CALL | 0.6 | PBMC2 | -0.3 |
| KCL22 | 14.4 | NALM6 | 5.7 | PBMC13 | 9.8 |
| MV4-11 | 19.5 | REH | 0.8 | PBMC14 | 4.3 |
| THP1 | 7.9 | DND41 | 2.0 | CD34+ | 4.0 |
| SKNO1 | 5.8 | HSB2 | 1.3 | F107 | 1.2 |
| CMK | 22.8 | Loucy | 4.2 | F188 | 0.9 |
| KASUMI | 11.6 | RPMI8402 | 18.3 | MSC | 4.1 |
| NB4 | 24.1 | PDX B-ALL1 | 30.5 | | |
| CMY | 18.3 | PDX B-ALL2 | 1.8 | | |
| HEL | 32.5 | PDX B-ALL3 | 14.8 | | |
| NOMO1 | 11.3 | PDX B-ALL4 | 12.8 | | |
| PDX AML1 | 4.3 | PDX B-ALL5 | 4.7 | | |
| PDX AML2 | 8.4 | PDX B-ALL6 | 6.6 | | |
| PDX AML3 | 5.3 | PDX T-ALL1 | 3.1 | | |
| PDX AML4 | 8.7 | PDX T-ALL2 | -1.0 | | |
| PDX AML5 | 1.9 | PDX T-ALL3 | 4.1 | | |
| PDX AML6 | 0.8 | PDX T-ALL4 | 4.2 | | |
| PDX AML7 | 7.3 | PDX T-ALL5 | 9.0 | | |
| PDX CML1 | 3.9 | PDX T-ALL6 | 2.7 | | |

E

**Ricolinostat x Clofarabine ZIP area**

| Myeloid Leukemia | | Lymphoid leukemia | | Healthy CTR | |
|---|---|---|---|---|---|
| Name | ZIP | Name | ZIP | Name | ZIP |
| HL60 | 18.6 | HAL01 | 19.3 | PBMC1 | 3.6 |
| K562 | 41.6 | MHH-CALL | 4.3 | PBMC2 | 8.1 |
| KCL22 | 8.2 | NALM6 | 9.4 | PBMC3 | 6.9 |
| MV4-11 | 21.1 | REH | 2.7 | PBMC4 | 9.2 |
| THP1 | 6.8 | DND41 | 7.3 | CD34+ | 8.8 |
| SKNO1 | 7.2 | HSB2 | 5.0 | F107 | 1.0 |
| CMK | 25.2 | Loucy | 7.8 | F188 | 7.4 |
| KASUMI | 12.3 | RPMI8402 | 18.1 | MSC | 3.8 |
| NB4 | 29.5 | PDX B-ALL1 | 17.3 | | |
| CMY | 16.2 | PDX B-ALL2 | 8.0 | | |
| HEL | 35.9 | PDX B-ALL3 | 8.1 | | |
| NOMO1 | 20.7 | PDX B-ALL4 | 5.8 | | |
| PDX AML1 | 15.9 | PDX B-ALL5 | 2.3 | | |
| PDX AML2 | 12.4 | PDX B-ALL6 | 5.3 | | |
| PDX AML3 | 17.3 | PDX T-ALL1 | 3.7 | | |
| PDX AML4 | 8.5 | PDX T-ALL2 | 0.0 | | |
| PDX AML5 | 7.6 | PDX T-ALL3 | 13.5 | | |
| PDX AML6 | 10.3 | PDX T-ALL4 | 4.4 | | |
| PDX AML7 | 18.2 | PDX T-ALL5 | 15.6 | | |
| PDX CML1 | 7.5 | PDX T-ALL6 | -0.3 | | |

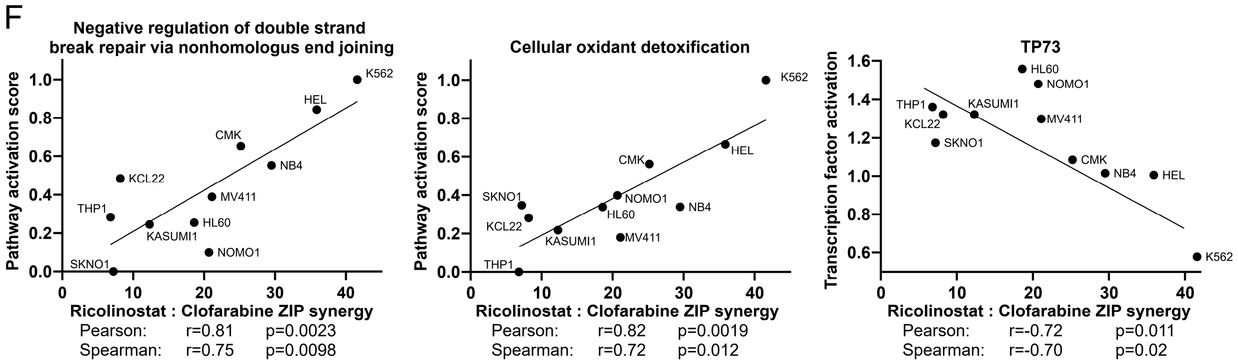


**Supplementary Figure 6. Targeting HDAC6 sensitizes myeloid leukemia cells towards standard chemotherapeutics Cytarabine and Clofarabine.** (A) Proliferation curve determined by trypan blue assay of K562 control (CTR) and K562 HDAC6-KO clone C2 treated with 2 µM Cytarabine over the course of 96 h (unpaired t-test, ns= not significant, n=3). (B) Bar plot of the fold change of the Caspase 3/7 activity of K562 CTR and K562 HDAC6-KO C2 treated with 2 µM Clofarabine over the course of 72h (unpaired t-test, *p< 0.05, **p< 0.01, ****p< 0.0001, n=3). (C) Bar plot showing a cell cycle analysis of K562 HDAC6-KO cells treated with Cytarabine (unpaired t-test, ns= not significant, *p< 0.05, **p< 0.01, ***p< 0.001, ****p< 0.0001, n=3). (D-E) Tables summarizing the most synergistic area ZIP scores for the combination of Ricolinostat with Cytarabine (D) or Clofarabine (E) across various leukemia subtypes. (F) Pathway and transcription factor activities were inferred from gene expression data of AML and CML cell lines and correlated with ZIP synergy scores of the Ricolinostat and Clofarabine combination (Pearson & Spearman correlation).

### Supplementary Table 1: Compounds List in the Drug screening Library

| Inhbitor | Mechanism of action / Target | Supplier | Reference |
|---|---|---|---|
| 5-Azacytidine | Antimetabolites | Selleckchem | S1782 |
| 6-Mercaptopurine | Antimetabolites | Selleckchem | S4504 |
| 6-Thioguanine | Antimetabolites | Selleckchem | S1774 |
| Clofarabine | Antimetabolites | Selleckchem | S1218 |
| Cytarabine | Antimetabolites | Selleckchem | S5582 |
| Cyclocytidine HCL | Antimetabolites | Selleckchem | S1973 |
| Nelarabine | Antimetabolites | Selleckchem | S1213 |
| Methotrexate | Antimetabolites | Selleckchem | S1210 |
| Vinblastine | Antimitotics | Selleckchem | S4505 |
| Vincristine | Antimitotics | Selleckchem | S1241 |
| Mitoxantrone | Topoisomerase | Selleckchem | S2485 |
| Daunorubicin | Topoisomerase | Selleckchem | S3035 |
| Amsacrine | Topoisomerase | Selleckchem | S5627 |
| Dexamethasone | GC/GCR complex | Selleckchem | S1322 |
| Prednisolone | GC/GCR complex | Selleckchem | S2570 |
| Belinostat | Histone deacetylases | Selleckchem | S1085 |
| CI-994 (Tacedinaline) | Histone deacetylases | Selleckchem | S2818 |
| Panobinostat | Histone deacetylases | Selleckchem | S1030 |
| Romidepsin | Histone deacetylases | Selleckchem | S3020 |
| Entinostat | Histone deacetylases | Selleckchem | S1053 |
| Ricolinostat | Histone deacetylases | Selleckchem | S8001 |
| GSK343 | Histone methyltransferase | Selleckchem | S7164 |
| Bortezomib | Proteasome | Selleckchem | S1013 |
| MLN-9708 | Proteasome | Selleckchem | S2181 |
| Ganetespib | Heat shock protein 90 | Selleckchem | S1159 |
| PUH71 | Heat shock protein 90 | Selleckchem | S8039 |
| AUY922 (LUMINESPIB) | Heat shock protein 90 | Selleckchem | S1069 |
| Alisertib | Aurora Kinase | Selleckchem | S1133 |
| Aurora A Inhibitor I | Aurora Kinase | Selleckchem | S1451 |
| Barasertib | Aurora Kinase | Selleckchem | S1147 |
| Volasertib | Polo like kinases | Selleckchem | S2235 |
| BI2536 | Polo like kinases | Selleckchem | S1109 |
| LY2835219 | Cyclin dependent kinases | Selleckchem | S5716 |
| Dinaciclib | Cyclin dependent kinases | Selleckchem | S2768 |
| SY-1365-THZ1 | Cyclin dependent kinases | Selleckchem | S7549 |
| Axitinib | Tyrosine kinases | Selleckchem | S1005 |
| Midostaurin | Tyrosine kinases | Selleckchem | S8064 |
| Dovitinib | Tyrosine kinases | Selleckchem | S7765 |
| Pexidartinib | Tyrosine kinases | Selleckchem | S7818 |
| Lestaurtinib | Tyrosine kinases | Biomol | LKT-L1875 |
| Quizartinib | Tyrosine kinases | Selleckchem | S1526 |
| Decitabine | Antimetabolites | Selleckchem | S1200 |
| Pacritinib | Tyrosine kinases | Selleckchem | S8057 |

| | | | |
|---|---|---|---|
| Idelalisib | Tyrosine kinases | Selleckchem | S2226 |
| Dactolisib (BEZ235) | Tyrosine kinases | Selleckchem | S1009 |
| Ro 08-2750 | Tyrosine kinases | Medchem | HY-108466 |
| ARQ-092 (Miransertib) | Tyrosine kinases | Selleckchem | S8339 |
| Everolimus | mTOR | Selleckchem | S1120 |
| Temsirolimus | mTOR | Selleckchem | S1044 |
| AT9283 | Tyrosine kinases | Selleckchem | S1134 |
| CYT387 (Momelotinib) | Tyrosine kinases | Selleckchem | S2219 |
| Ruxolitinib | Tyrosine kinases | Selleckchem | S1378 |
| Fedratinib (TG101348) | Tyrosine kinases | Selleckchem | S2736 |
| Tipifarnib | Farnesyltransferase | Selleckchem | S1453 |
| Lonafarnib | Farnesyltransferase | Selleckchem | S2797 |
| Sorafenib (Tosylate) | Tyrosine kinases | Selleckchem | S1040 |
| Regorafenib | Tyrosine kinases | Selleckchem | S5077 |
| Cobimetinib | Mitogen activated protein kinases | Selleckchem | S8041 |
| MEK162 (Binimetinib) | Mitogen activated protein kinases | Selleckchem | S7007 |
| Trametinib | Mitogen activated protein kinases | Selleckchem | S2673 |
| Tirabrutinib | Bruton’s tyrosine kinase | Medchem | HY-15771 |
| Ibrutinib | Bruton’s tyrosine kinase | Medchem | S2680 |
| Imatinib | Tyrosine kinases | Medchem | S1026 |
| Ponatinib | Tyrosine kinases | Medchem | S1490 |
| Bosutinib | Tyrosine kinases | Medchem | S1014 |
| Dasatinib | Tyrosine kinases | Medchem | S5254 |
| Staurosporin | Tyrosine kinases | Medchem | S1421 |
| BSI-201 (Iniparib) | Poly(ADP-ribose)-Polymerasen | Medchem | S1087 |
| Olaparib | Poly(ADP-ribose)-Polymerasen | Medchem | S1060 |
| ABT-199 (Venetoclax) | BCL2 apoptosis regulators | Medchem | S8048 |
| Obatoclax | BCL2 apoptosis regulators | Medchem | S1057 |
| QNZ (EVP4593) | NF-κB | Medchem | S4902 |
| Omaveloxolone | Inflmmation modulation | Medchem | S7672 |
| Birinapant | Cellular inhibitors of apoptosis | Medchem | S7015 |
| Selinexor | Exportin | Medchem | S7252 |
| AZD6738 | Tyrosine kinases | Medchem | S7693 |
| Homoharringtonine | Protein translation | Medchem | S9015 |
| Bexarotene | Retinoid X recepetors | Medchem | S2098 |
| Nintedanib (BIBF1120) | Tyrosine kinases | Medchem | S1010 |
| BAY 80-6946 (Copanlisib) | Tyrosine kinases | Medchem | S2802 |
| Palbociclib | Cyclin dependent kinases | Medchem | S1116 |
| Birabresib | BET bromodomain | Medchem | S7360 |
| Tegaserod Maleate | 5-HT4 receptor | Medchem | S5401 |
| 5-nonyloxy tryptamine | Serotonin receptor | Sigma Alderich | 157798-13-5 |

#### Supplementary Table 2: Clinical characteristics of the leukemia patients xenografted in NSG mice

*Related to Figure 2, 3, 4 and 6*

| PDXs | Samples used for xenograft | Age at first diagnosis (yrs) | Sex | FAB (AML) | Karyotype | Genetic lesions |
|---|---|---|---|---|---|---|
| CML | initial | 14,9 | F | - | 46XX, t(9:22)(q34,q11); 47, idem, +21 | BCR::ABL1, trisomy 21, trisomy 20 |
| AML1 | relapse | 15,3 | M | M0 | 46, XY | RARA, RUNX1 |
| AML2 | initial | 14,3 | M | M5 | 46, XY | FLT3-ITD |
| AML3 | relapse | 51,0 | M | - | Normal Kryotype | FLT3-mut, NPM1 |
| AML4 | relapse | 0,5 | F | M0 | 46, XX, inv(9)(p11q12) | MLL::AFF1 , NRAS |
| AML5 | relapse | 71 | M | - | Normal Karyotpe | ASXL1, DNMT3A, FLT3-ITD, FLT3-TKD, NPM1, TET2 |
| AML6 | initial | 9,6 | M | M4 | 46, XY, inv(16)(p13q24) | CBFB::MYH11, FLT3-TKD, NRAS, PTPN11 |
| AML7 | relapse | 47,0 | F | M4 | 46,XX, ins (10;11) (p12;q23q23) | KMT2A::AF10, KRAS:p.G12A, BCOR |
| B-ALL1 | relapse | 12,1 | F | - | - | TCF3::PBX1 |
| B-ALL2 | relapse | 4,5 | M | - | 46, XY, t(9;22)(q34;q11) | BCR::ABL1, IKZF1 |
| B-ALL3 | relapse | 11,0 | M | - | arr[GRCh37] (Y)x0,(2-4)x1,5q23.2q31.3(125944131_142444598)x1, 5q31.3q32(142804687_148958726)x1,(7,12,13,15-17)x1,(20)x1[0.9] | TP53 NM_000546.5:c.844C>T, p.Arg282Trp |
| B-ALL4 | initial | 3,1 | M | - | SNP-array derived karyotype: 54,XY,+X,+Y,CN-LOH(1-6),del(6)(p22.2)/HIST1x2,CN-LOH(7,8),+9,+9,CN-LOH(10-13),+14,+14,CN-LOH(15-20),+21,+21, CN-LOH(22); | Mono-clonal non classical-high hyperdiploid ("masked near-haploid") |
| B-ALL5 | relapse | 5,4 | M | - | SNP-array derived karyotype: 53,XY,+X,+Y,CN-LOH(1-6),del(6)(p22.2)/HIST1x2,CN-LOH(7,8),+9,CN-LOH(10-13),+14,+14,CN-LOH(15-19),del(19)(p13.3),CN-LOH(20),+21,+21,CN-LOH(22) | Mono-clonal non classical-high hyperdiploid ("masked near-haploid") |
| B-ALL6 | initial | 15,0 | F | - | Karyotype by G-banding: 46,XX[10]; karyotype by SNP-array: 50,XX,+X,+X,CN-LOH(1-20,22),+21,+21 | Mono-clonal non classical-high hyperdiploid ("masked near-haploid") |
| B-ALL7 | relapse | 7,0 | M | - | 29, XY | TP53 NM_000546.5:c.846_847insGCCGG,p.Arg283AlafsTer64 |
| T-ALL1 | initial | 8,1 | F | - | - | - |
| T-ALL2 | initial | 3,3 | M | - | - | - |
| T-ALL3 | initial | 7,7 | F | - | - | CDKN2A |
| T-ALL4 | initial | 1,5 | F | - | - | Cornelia de Lange Syndrome |
| T-ALL5 | initial | 4,5 | F | - | - | - |
| T-ALL6 | initial | 14,3 | F | - | 47, XX, t(5;14)(q35.1;q32.2), +8 | IKZF1, trisomy 8 |

#### 2. Supplemental Material and Methods:

**CRISPR-Cas9 mediated knockout (KO)**: K562 wild type cells were transduced with a CAS9 expression plasmid (Addgene, Ref: 108100) via Lenti-X Packaging Single Shots (Takara, Ref: 631276) using HEK293T cells. gRNAs were cloned into the lentiviral expression plasmid with GFP or mCherry (Addgene, Ref: 108098 or 108099). Virus production was was done in Lenti-X HEK293T cells using Lenti-X Packaging

Single Shots (Takara). After 72 h in total, the lentiviral supernatant was harvested, filtered through a 0.45 µm filter and added to constitutively expressing Cas9 K562 cells (0.4 x $10^6$ cells) that were previously generated. After transduction, positive cells were selected by the fluorescent marker, GFP or mCherry.[1]

*Generation of KCL-22, MV4-11, NALM6, and C1498 HDAC6-KO cells:* The transfection was carried out using the Amaxa Nucleofection system SF Cell Line Kit (Lonza, Ref: V4XC-2032). For 2 x $10^5$ cells, 100 pmol of Cas9-GFP protein (IDT, Ref: 10008100) was mixed with 120 pmol of gRNA (crRNA:tracrRNA 1:1) and assembled for 20 min at RT. Afterwards, the labelled ssODN was added and the mixture was combined with the cell suspension (resuspended cells in Nucleofector solution SF) and the electroporation enhancer (IDT). The complete volume was transferred to the Nucleocuvette module, placed in the 4D-Nucleofector system (Lonza) and electroporated with the program CA-137 for KCL22, DJ-100 for MV4-11, CV-104 for NALM6 and CM-138 for C1498. After 24h, the cells were sorted for GFP and monoclonal selected via semi-solid cloning. The following guide RNA (gRNA) sequences were utilized:

| Construct | Sequence (5' – 3') |
|---|---|
| gRNA_HDAC6 human 1 | GCA CAT CCC AAT CTA CGA TA |
| gRNA_HDAC6 human 2 | CTA TTG CAT GTT CAA CCA CG |
| gRNA_HDAC6 murine 1 | GAA CAT CCC AAT CCA CGA TTA GG |
| gRNA_HDAC6 murine 2 | ACA CGT ATA ATA CAC TGC CAG GG |

**ShRNA mediated HDAC6-knockdown (KD):** SMARTvector lentiviral shRNA constructs (Horizon Discovery) containing a human EF1α promoter and TurboRFP fluorescent reporter were used to knock down HDAC6 expression, following the manufacturer's instructions. The specific shRNA sequences used are listed below.

| Target | Mature antisense sequence (5' – 3') | Cat. number |
|---|---|---|
| sh1 | CAC ATG ATC CGC AAG ATG C | V3SVHS09-10351093 |
| sh2 | CCA CGT ACT CAG CAC TGT G | V3SVHS09-8678884 |
| sh3 | ATC CCA ATC CAC AAT CAG G | V3SVHS09-7082476 |
| Non-targeting (NT) control | | SVC17012404 (hEF1a-Turbo RFP) |

**HDAC6-knockin (KI):** For the HDAC6 knock-in experiments, the HDAC6 plasmid (Addgene, Ref: 13823) was subcloned into a pCDH-EF1α-GFP lentiviral vector (System Biosciences). K562 HDAC6-KO cells were transduced with the lentiviral construct, and GFP-positive cells were FACS-sorted.

**qPCR.** For RNA extraction, cell pellets were first lysed using QIAzol Lysis Reagent (QIAGEN), followed by isolation with the Maxwell® RSC Viral Total Nucleic Acid Purification Kit (Promega) using the Maxwell® RSC 48 instrument. A total of 2 µg of purified RNA from each sample was used as a template for cDNA synthesis with the QuantiTect Reverse Transcription Kit (QIAGEN). Quantitative PCR (qPCR) was performed using a real-time PCR system (Bio-Rad). The mean Ct values of the housekeeping genes B2M and GAPDH were used as internal controls to normalize target gene expression levels. Human primer sequences are listed below:

| Target | Forward primer sequence (5'-3') | Reverse primer sequence (5'-3') |
|---|---|---|
| **RNAseT2** | GGC ATA CTG GCC TGA CGT AAT | CTT TTC CCA CTC ATG CTT CCA GA |
| **B2M** | GTA TGC CTG CCG TGT GAA C | AAA GCA AGC AAG CAG AAT TTG G |
| **β-actin** | GCA CTC TTC CAG CCT TCC | CTC GAA GCA TTT GCG GTG |

**Fluorescence microscopy:** Microscope slide cover slips were coated with poly-D-lysine (Gibco, Ref: A3890401) for 1 hour, washed with PBS, and seeded with treated or genetically engineered cells for adhesion. Cells were fixed in 4% formaldehyde (Merck, Ref: 1.00496), quenched in Tris-buffered saline, and permeabilized with 0.01% Triton X (Roth, Ref: 3051.2).[2] Blocking was performed overnight with 10% BSA at 4°C. Primary and secondary antibodies were used per supplier recommendations (see supplementary) with 2-hour incubations, followed by washes. Nuclei were stained with DAPI (Thermo Fisher, Ref: 62248), and slides were mounted with ProLong™ Gold (Invitrogen, Ref: P10144), dried for 48 hours, and imaged using an Axio Observer and Apotome 3 (Zeiss). After equalization via the histogram the signal of single cells and the background was quantified in ImageJ. After subtraction of the background the signal was normalized to the controls and the significance calculated in an unpaired t test (n=20).

**ATAC-seq**: The data was uploaded and analyzed using the galaxy platform. After an initial quality control with FastQC the adapters were trimmed with Cutadapt. The mapping was done via Bowtie2 and filtered afterwards for mapping quality (>= 30), proper pairing and chromosomal origin (!= mitochondrial), while PCR amplified duplicates were flagged via MarkDuplicates. Subsequently the files underwent a conversion from BAM to BED and were used with MACS2 for peak calling. The given BedGraph files were then converted to the BigWig format and scaled to 1.000.000/number of reads. For the generation of the heatmap the bigwig files were averaged trough BigWig average, given to computeMatrix and finally visualized with plotHeatmap tools. Track plots were generate with the help of the pyGenomeTracks package. Differential gene expression analysis was done via DiffBind. The differentially open genes were processed similar to the RNAseq & Proteomics data. The cut-off for the overlap analysis was FDR<0.05. As reference genome hg38 was used.

**T cell co-culture activation assay:** C57BL/6J 6-8 weeks old mice were infected I.V. with 5 x $10^5$ PFU LCMV Armstrong strain.[3] After 2 weeks mice were sacrificed and splenic pan T cells were isolated with Pan T Cell Isolation Kit (Miltenyi Biotec, 130-095-130) according to manufacturer instructions. After determination of the density via trypan blue counting the cells were resolved in culture media with 100 U/ml IL2 (Gibco, Ref: 00-02-250) and with a concentration of 1 *10^6 cells per ml. 0.05*10^6 C1498 cells were treated 24 h prior to the co-culture experiment with 1 µM of Ricolinostat. Afterwards the treatment media was discarded and the cells co-incubated with 0.1 *10^6 of the isolated T cells. For the killing curve cells were diluted in three 1:1 steps. Separately 1*10^6 T cells were incubated with a 1:1 ratio of supernatant from treated C1498 to fresh media. The supernatant was filtered through a 0.45 µm strainer to exclude the cells. Also 1*10^6 T cells were treated with 1 µM of Ricolinostat. After 24h the cells were either analyzed in the CytoFLEX to determine the viability via DAPI (Invitrogen, 62248) or fixed for intracellular staining. After 24 hours cells were stained for surface markers, next intracellular cytokines staining performed with Foxp3 / Transcription Factor Staining Buffer Set (Invitrogen, 00-5523-00) according to manufacturer instructions with Fixable Viability Dye (Invitrogen, 65-0863-14,). Flow cytrometry was performed on Cytoflex (Beckman Coulter) and data analyzed with FlowJo_v10.7.2 software. For each of the 8 mice duplicates were used. Significance was determined via unpaired t test (n=16).

**NK-cell killing assay**: PBMCs were isolated from healthy donors' buffy coats, provided by the blood bank of the University Hospital Düsseldorf, using Ficoll-Paque TM Plus GE (Sigma-Aldrich, Ref: GE17-1440-02) density gradient centrifugation. After red blood lysis, samples were frozen with a cryopreservation medium containing FBS with 10% DMSO (Sigma-Aldrich, St. Louis, USA). For the experiments, PBMCs were thawed in RPMI GlutaMAX medium with 10% human serum (Sigma-Aldrich, Ref: H4522), 200 U/ml IL-2 (Miltenyi Biotec, Ref. 130-097-743) and 0.3 ng/ml IL-15 (Miltenyi Biotec, Ref. 130-095-760) and incubated at 37°C with 5% CO2 overnight. Target cells were harvested and stained with CFSE (Thermo Fisher Scientific, Ref. C34570) and seeded into a 96-well U-bottom with 1*10^5 cells per well. PBMCs were harvested and seeded into the same plate with 1*10^6 cells per well. Target cells and PBMCs were co-cultured to test the NK cell activity. Mono-cultures of the respective cells were used as controls to be able to calculate the excess killing and death rates. After 4 hours of culture, cells were stained for flow cytometry.

**Murine C57BL/6J experiments:** Mice were sacrificed and spleen, lymph node (LN) and bone marrow (BM) isolated. Cells were stimulated for 1 hour with 5 ng/ml PMA (Sigma-Aldrich, Ref: P1585) and 25 ng/ml Ionomycin (Invitrogen, Ref: I24222) for cytokine accumulation, next Brefeldin A (Merck, Ref: B5936) was added to the culture and cells were incubated for additional 5 hours. Next, cells were stained for surface markers and intracellular cytokines staining (see supplementary) performed with Foxp3 / Transcription Factor Staining Buffer Set (Thermo Fisher Scientific, Ref. 00-5523-00) with Fixable Viability Dye (Invitrogen, Ref: 65-0863-14).[3]

Another cohort of mice was used to further investigate the antitumor efficacy of HDAC6i treatment. The engraftment of leukemia cells was confirmed 7 days post tumor inoculation via IVIS and divided randomly to different treatment groups. In total three cycles of treatments (successive 5-day daily treatment course followed by one day break) with either vehicle control (10% DMSO+ 40% PEG300+ 5% TWEEN80 + 45% ddH2O) or 50mg/kg Ricolinostat (Selleckchem) were administered i.p.. The tumor burden of each mouse was assessed by monitoring the bioluminescent signals and quantifying the region of interest (ROI) weekly. Health conditions and body weight from all animals were closely monitored throughout the experimental period. After the endpoint of the experiment, a survival analysis was performed.

**Western Blotting:** Cells were harvested by centrifugation at 400 x g for 5 min at 4°C and washed three times with ice-cold PBS and then snap-frozen in liquid nitrogen.[1] 100-300 µL protein lysis (see below) buffer and was used for 5 million cells. Cells were lysed on ice for 30 minutes with periodically vortexing, centrifuged two times (10000 x g for 20 min at 4 °C) and protein quantification was performed by BCA-Assay (Thermo Fisher Scientific, #23227). 10 - 20 µg lysate were separated by SDS-PAGE at 50 V for 30 mins during stacking phase with subsequent 100 V for 2h during separation phase and blotted onto 0.45 µm nitrocellulose membrane (Cytiva, #10600002) at 100 V for 1h or overnight (30 V, 16 h). Membranes were washed twice with TBS and analyzed for their protein content by ponceau staining (Sigma-Aldrich #P7170) staining. After three washes with TBS-T, membranes were blocked in 5% BSA (Sigma-Aldrich, #A3294) TBS-T solution, washed three times with TBS-T and incubated with primary antibody solution in 5% BSA solution overnight at 4°C. Membranes were washed three times with TBS-T, incubated for 1h with secondary HRP-conjugate (Cell Signaling Technologies, #7074 and #7076) at 1:2000 in TBS-T solution, washed again three times in TBS-T and lastly one time in TBS. For visualization ECL-Solution (Cytiva, #GERPN2106) was used as per manufacturer's instruction and image was captured using JESS.

***List of antibodies used in immunoblots***:

| Target | Dilution | Origin | Supplier | Reference |
|---|---|---|---|---|
| **HDAC6** | 1:1000 | Rabbit | Cell signaling | 7558S, 7612 |
| **ac-α-Tubulin** | 1:1000 | Rabbit | Cell signaling | 5335S |
| **LAMP1** | 1:1000 | Rabbit | Cell signaling | 9091T |
| **RNAse T2** | 1:1000 | Rabbit | Abcam | ab 140191 |
| **RNAse T2** | 1: 500 | Mouse | Santa Cruz | sc-393729 |
| **PARP** | 1:1000 | Rabbit | Cell signaling | 9542S |
| **GAPDH** | 1:1000 | Rabbit | Cell signaling | 5174S |
| **β-Actin** | 1:1000 | Mouse | Cell signaling | 12262S |
| **Nucleolin** | 1:1000 | Rabbit | Cell signaling | 14574S |
| **α-Tubulin** | 1:1000 | Rabbit | Cell signaling | 2144S |
| **Anti-rabbit** | 1:2000 | Goat | Cell signaling | 7074S |
| **Anti-mouse** | 1:2000 | Horse | Cell signaling | 7076S |

***List of antibodies used in Fluorescence microscopy***:

| Target | Dilution | Origin | Supplier | Reference |
|---|---|---|---|---|
| **LAMP1** | 1:50 | Rat | Santa Cruz | 19992 |
| **RNAse T2** | 1:50 | Mouse | Santa Cruz | 393729 |
| **Anti-Mouse IgG Alexa Fluor™ plus 488** | 1:1000 | Mouse | Invitrogen | A32723TR |
| **Anti-Rat Alexa Fluor™ plus 647** | 1:1000 | Rat | Invitrogen | A78947 |

***List of antibodies used in FACS***:

| Target | Origin | Supplier | Reference |
|---|---|---|---|
| **CD3** | Rat | Invitrogen | 25-0038-42 |
| **CD56** | Mouse | BD Pharmingen | 557711 |
| **CD107a** | Mouse | BioLegend | 328630 |
| **PD-L1** | Mouse | BioLegend | 329732 |
| **ULBP2/5/6** | Mouse | BD OptiBuild | 748131 |
| **CD80** | Mouse | BD Horizon | 567426 |
| **B2M** | Mouse | BD | 656645 |
| **HLA-ABC** | Mouse | Invitrogen | 12-9983-42 |
| **CD3** | Hamster | eBioscience | 47-0031-82 |
| **CD8** | Rat | BD Biosciences | 563068 |
| **CD4** | Rat | Invitrogen | 63-0042-82 |
| **KLRG1** | Hamster | BD Biosciences | 740553 |
| **GZMB** | Rat | eBioscience | 12-8898-82 |
| **IL2** | Rat | eBioscience | 45-7021-82 |
| **IFN-γ** | Rat | eBioscience | 17-7311-82 |
| **TNFα** | Rat | eBioscience | 48-7321-82 |
| **CD107a** | Rat | BD Biosciences | 560647 |
| **IL7R/CD127** | Rat | eBioscience | 45-1271-82 |
